# Fertilization by short-term stored sperm alters DNA methylation patterns in common carp (*Cyprinus carpio*) embryos at single-base resolution

**DOI:** 10.1101/2024.02.18.580899

**Authors:** Yu Cheng, Songpei Zhang, Rigolin Nayak, Pavlína Vechtová, Fabian Schumacher, Pavla Linhartová, Ievgeniia Gazo, Zuzana Linhartová, Swapnil Gorakh Waghmare, Burkhard Kleuser, Abhipsha Dey, Vladimíra Rodinová, Marek Rodina, Jan Sterba, Sayyed Mohammad Hadi Alavi, Catherine Labbé, Otomar Linhart

**Author notes:** **Corresponding author:** prof. Ing. Otomar Linhart, DrSc, University of South Bohemia in České Budějovice, Faculty of Fisheries and Protection of Waters, South Bohemian Research Center of Aquaculture and Biodiversity of Hydrocenoses, Zátiší 728/II, 389 25 Vodňany, Czech Republic.

## Abstract

Short-term storage of sperm *in vitro* is widely used for artificial fertilization in aquaculture. It has been shown that short-term storage affects sperm motility characteristics, resulting in diminished fertility. However, the detrimental effects of short-term sperm storage on embryos development have remained largely unexplored in single-base methylome resolution. The main aim of the present study was to investigate DNA methylation in the offspring of common carp (*Cyprinus carpio*) derived from short-term stored sperm. Sperm were stored in artificial seminal plasma on ice (0-2^°^C) for 0, 3 and 6 days *in vitro*, fertilization was performed using oocytes from a single female, and embryos were collected at the mid-blastula stage. Sperm and embryo DNA was extracted, analysed by liquid chromatography with tandem mass spectrometry (LC-MS/MS) and whole genome bisulfite sequencing (WGBS). DNA methylation was then assessed. Sperm storage showed negative effects on motility, viability and DNA integrity, but had no effect on global DNA methylation of spermatozoa and resulting embryos. Results from the WGBS showed that methylation of 3313 genes was affected in the embryos fertilized with the 6-day-stored sperm, and the differentially methylated regions (DMRs) identified were mainly involved in cell adhesion, calcium, mitogen-activated protein kinase and adrenergic signalling, melanogenesis, metabolism and RNA transport. Such results suggest that prolongation of storage time may have certain impacts on embryonic development. These initial results provide valuable information for future consideration of the DNA methylome in embryos generated from short-term stored sperm, which is widely used for genetic management of broodstock in aquaculture.

## Introduction

Information not contained in a DNA sequence can be transmitted from parents to offspring through gametes. This is known as epigenetic inheritance, in which non-DNA sequence-based epigenetic information can be inherited across several generations (Daxinger and Whitelaw 2012; Fitz-James and Cavalli 2022). DNA methylation is a widely studied heritable epigenetic modification that occurs predominantly in the CpG context in animals and plays a crucial role in regulating gene expression by recruiting proteins involved in gene repression and inhibiting transcription factors without altering the DNA sequence (Egger et al. 2004; Ohka et al. 2011). In reproduction, a number of environmental factors or physiological conditions can cause abnormal DNA methylation in gametes (Zhang et al. 2023b). For example, short-term and long-term storage of sperm can lead to changes in sperm DNA methylation (Shazada et al. 2023; Zhang et al. 2023b). In this context, the DNA methylation of spermatozoa is strongly influenced by the use of an extender for short-term storage and the type of cryoprotectant used for cryopreservation. Cheng et al. (2023) reported that global DNA methylation changes induced by short-term-storage in common carp (*Cyprinus carpio*) spermatozoa can be prevented if sperm storage was carried out in an extender. Depincé et al. (2020) stated that dimethylsulfoxide and 1,2-propanediol decreased global DNA methylation in goldfish (*Carassius auratus*) spermatozoa, but methanol, as an effective cryoprotectant, did not cause any notable changes. Interestingly, epigenetic modifications in gametes under environmental and/or physiological conditions show species specificity. For instance, in contrast to goldfish sperm, global DNA methylation in zebrafish (*Danio rerio*) sperm increased after cryopreservation using methanol as a cryoprotectant (Depincé et al. 2020). In addition, it is worth paying attention to epigenetic reprogramming in embryos, because of its crucial role in embryonic development (Feng et al. 2010). It has been reported that sperm preserved at –20^°^C without cryoprotectants is epigenetically safe to use for intracytoplasmic sperm injection, as the dynamic epigenetic reprogramming of embryos derived from frozen sperm was similar to that of embryos derived from fresh sperm (Chao et al. 2012). However, epigenetic changes in response to the environmental stressor mediate an organism’s adaptation to the environment. In shortfin molly (*Poecilia mexicana*), the acclimatised epigenome under hydrogen sulfide could be inherited by subsequent generations, facilitating the adaptation of *P. mexicana* to sulfidic environments (Kelley et al. 2021).

In aquaculture, short-term storage of sperm is a convenient and valuable method of preserving sperm under refrigerated conditions for artificial fertilization (Shazada et al. 2023). It is most beneficial to reduce the hatchery workload on the same day that females are stripped, and to conserve male sperm if females are delayed in maturing. However, spermatozoa are exposed to environmental stressors, such as reactive oxygen species (ROS) and natural aging during the storage (Dietrich et al. 2021). Importantly, oxidative stress in sperm may influence epigenetic reprogramming in early embryonic development (Wyck et al. 2018). Spermatozoa have compact nuclei, which distinguishes them from somatic cells. In addition, while DNA methylation remains stable in somatic tissues, its patterns and levels change dynamically during gametogenesis and embryonic development. Stability of sperm DNA methylation has been reported as being critical for developing embryos (Labbé et al. 2017; Donkin and Barrès 2018). In zebrafish, the paternal DNA methylation pattern is maintained throughout early embryonic development, but the maternal methylation pattern is reprogrammed to match the paternal model (Jiang et al. 2013; Murphy et al. 2018).

The literature reveals that environmental factors during gamete storage may affect the epigenome (Andreu-Noguera et al. 2023) and their epigenetic reprogramming is essential for embryonic development (Potok et al. 2013; Murphy et al. 2018). Sperm storage is mostly used traditionally in artificial reproduction of economically important species in aquaculture; compared to studies in model fish species, epigenetic research is more recent and scarcer (Liu et al. 2022). Common carp is one of the major finfish species in aquaculture, with global production increasing from 1205 thousand tonnes in 1990 to 4360 thousand tonnes in 2020 (Shazada et al. 2023). Offspring are produced by artificial reproduction, and short-term stored or cryopreserved sperm is widely used (Shazada et al. 2023). Therefore, it is worth assessing the safety of stored sperm for the offspring in terms of their gametes’ handling/storage and their application in artificial reproduction. In our recent study, sensitive methods based on liquid chromatography with tandem mass spectrometry (LC-MS/MS) were performed to evaluate the global epigenetic level (5-mdC) in sperm and embryos. Finally, there were no significant changes in epigenetic factors (Cheng et al. 2023). However, due to the high variability of global methylation level even with without significant difference observed between groups of embryos derived from fresh and stored sperm (Cheng et al. 2023), there may be different site-specific DNA methylation signatures (Khezri et al. 2019; He et al. 2022). More accurate techniques need to be developed that can provide higher resolution at the single base level, such as whole-genome bisulfite sequencing (WGBS), which is a valuable and extensively used approach. Moreover, understanding the epigenetic regulation of different features can help practical applications in aquaculture using different methodologies. For example, the quality of common carp gametes and embryos could be enhanced in the future by epigenetic modifications of the sperm, such as epigenomic editing and epigenetic selection (Metzger and Schulte 2016; Katayama and Andou 2021; Nakamura et al. 2021).

The aim of the present study was to investigate the sperm phenotypes and DNA methylation in the common carp embryos from artificial fertilization with short-term sperm storage. The fresh oocytes from one female were fertilized by stored sperm, and the differentially methylated regions (DMRs) and differentially methylated genes (DMGs) were identified in the embryos at the mid-blastula stage by direct deep sequencing to elucidate correlations between the DNA methylation profile in the embryos and the period of sperm storage.

## Materials and methods

### Animal ethics

The manipulations of animals in this study were performed in accordance with the relevant legal requirements and ethical standards. Specifically, the breeding and the supply of laboratory animals were authorised by the Ministry of Agriculture of the Czech Republic under reference numbers 56665/2016-MZE-17214 and 64155/2020-MZE-18134, while the use of laboratory animals was authorised under reference number 68763/2020-MZE-18134. In addition, the authors of this study, PL (CZ 03273), VR (CZ 02992), MR (CZ 01664) and OL (CZ 02815), hold a Certificate of Professional Competence according to Section 15d (3) of ACT No. 246/1992 Coll. on the Protection of Animals against Cruelty, which qualifies them to perform animal experiments.

### Broodstock and gamete collection

Five 3-year-old matured common carp males were chosen from broodstock with healthy and good-quality sperm. For each hormonal administration, a single intramuscular injection of carp pituitary (CP; 2 mg/kg b.w. dissolved in physiological solution) was performed to induce sperm maturation (Linhart et al. 2015). The milt of each male was collected with a syringe and stored on crushed ice (0–2 ^°^C) *in vitro* under aerobic conditions. At first, the quality of sperm was evaluated, and the best three sperm samples with motility ∼80% were selected for further experiments (Fig. 1). To induce final oocyte maturation, the female was treated with CP at 0.5 mg/kg b.w. (primary injection) and 2.7 mg/kg b.w. (secondary injection) with a 12 h interval. A good quality oocyte from a female (8.3 kg) in good physical condition was chosen for the fertilization test.

**Fig. 1.**
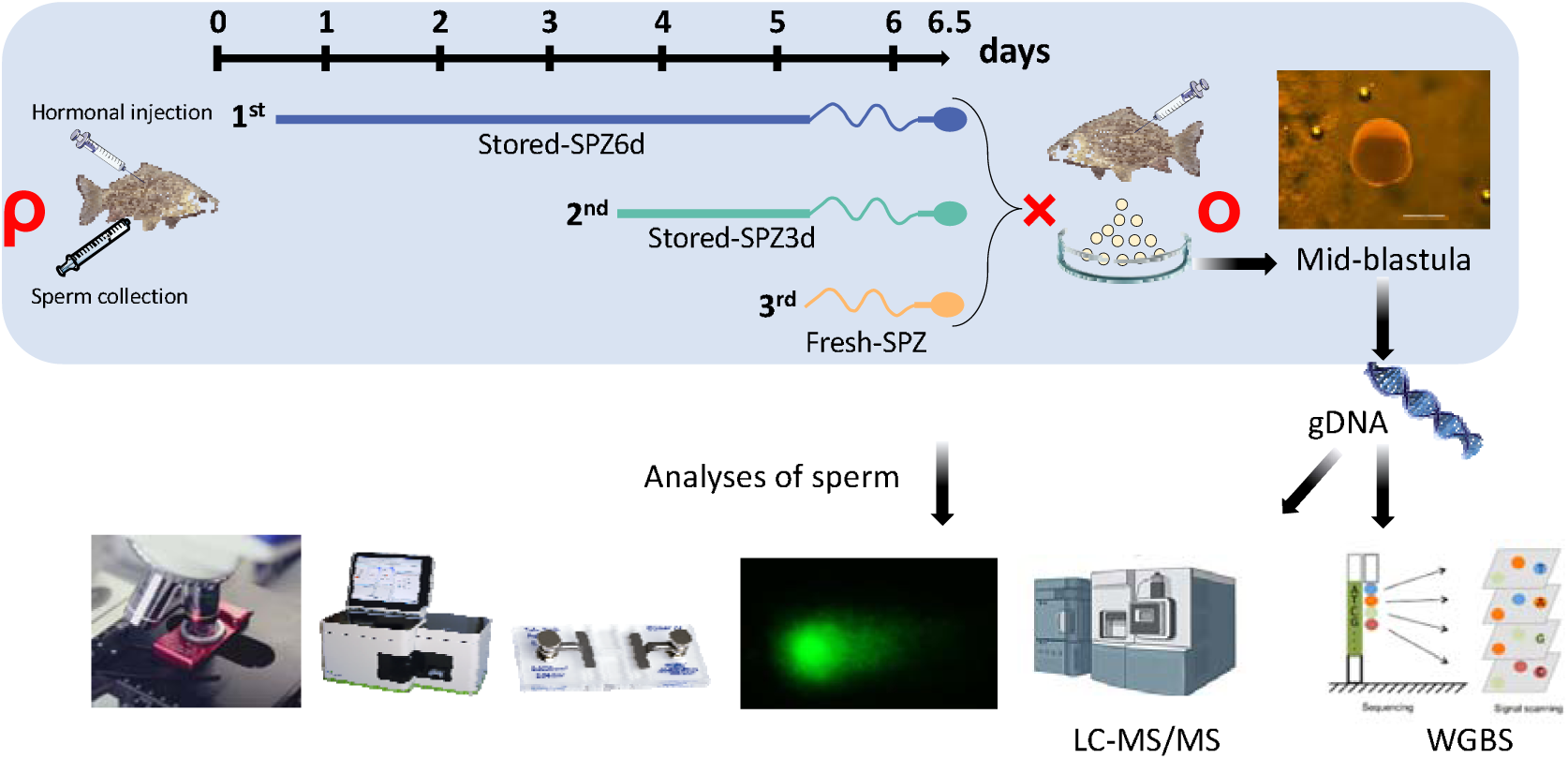
Experimental design and procedure for comparing fresh sperm (Fresh_SPZ) *vs*. stored sperm for 3 days (Store_SPZ3d) *vs*. stored sperm after 6 days (Store_SPZ6d) and their embryos. Three strippings were performed on each male (*n* = 3) at 12 h post each hormonal stimulation of spermiation, and sperm samples were immediately diluted with an extender and stored on ice (0–2 °C). The sperm from the first hormonal treatment was stored *in vitro* for 6 days *in vitro* (Stored_SPZ6d), the sperm from the second hormonal treatment for 3 days (Stored_SPZ3d), and the third stripped sperm was without storage *in vitro* (Fresh_SPZ). In total three sperm samples were used to fertilize eggs from one female on the same day. Sperm phenotypes (sperm kinetics, membrane integrity), DNA fragmentation and global DNA methylation (5-mdC/dC (%)) were analysed. DNA methylation of embryos in the single base resolution was sequenced by WGBS. A consistent colour scheme is used throughout the figures for stored and fresh sperm and the corresponding embryos.

### Experimental design for sperm storage *in vitro*

The experiment was designed to obtain fresh sperm and stored sperm *in vitro* from the same males to explore the effects of sperm storage on spermatozoa epigenetics and their next generation implications in common carp (Fig. 1). For each male, we stored sperm *in vitro* on ice after each stripping to make varied aged sperm for artificial fertilization and simultaneously to produce embryos with the same mother. Multiple hormone injections and male stripping were required to accomplish this, but it was done carefully to avoid harming the fish. In brief, three hormonal treatments were provided to the same three males at 3-day intervals (days 0, 3 and 6 of the experiment). Fresh sperm was collected at each 12 h post-hormonal treatment and immediately diluted with an extender (110 mM NaCl, 40 mM KCl, 2 mM CaCl_2_, 1 mM MgSO_4_, 20 mM Tris, pH 7.5 and 310 mOsmol/kg) (Cejko et al. 2018) at a ratio 1:1 (*v*:*v*). Fresh sperm (Fresh_SPZ), stored sperm for 3 days (Store_SPZ3d) and 6-day-stored sperm (Store_SPZ6d) were stored *in vitro* for 6-, 3-and 0-days post stripping (DPS), respectively, to achieve artificial fertilization at the same time. The Fresh_SPZ obtained from the third stripping was used as a control.

### Evaluation of sperm phenotypes

Spermatozoa motility, velocity and viability were assessed as phenotypic traits of sperm (Cheng et al. 2023). Spermatozoa motility was activated in distilled water containing 0.25% Pluronic F-127 at room temperature (21 ^°^C), and the percentage of motile spermatozoa and spermatozoa velocity were evaluated from recorded videos at 15 s post sperm activation. Spermatozoa membrane viability was assessed using the LIVE/DEAD Sperm Viability Kit (Invitrogen/Thermo Fisher Scientific Inc.). A flow cytometry with the S3e^TM^ Cell Sorter (Bio-Rad, Hercules, CA, USA) was used to estimate the ratio of live: dead spermatozoa.

### Assessment of sperm DNA integrity

The DNA integrity of sperm was evaluated using a modified alkaline single-cell gel electrophoresis (comet assay) as described previously (Gazo et al. 2022; Giassetti 2023). Briefly, microscope slides (OxiSelectST; Cell Biolabs, INC. USA) were pre-coated with 1% normal-melting-point agarose (Invitrogen) dissolved in TBE buffer (90 mM Tris base, 90 mM boric acid, 2.5 mM EDTA) 1 day before the comet assay. A volume of 0.6 µl of sperm was diluted with 2.5 ml of PBS (phosphate buffer solution). Diluted samples (100 µl) were mixed with 800 µl of agarose (low EEO, Sigma). Finally, 40 µl of this mixture was added to the slide and allowed to solidify at 4 ^°^C for 5 min. All slides were gently immersed in lysis buffer (2.5 M NaCl, 100 mM EDTA, 10 mM Tris, 1% Triton X-100, 10% DMSO, pH 10) and incubated overnight at 4 ^°^C. After the cells were lysed, the buffer was removed, and the slides were washed in TBE, and placed in a gel tank filled with electrophoresis buffer (TBE). Electrophoresis was performed for 15 min at 50 V. The slides were washed three times with distilled H_2_O following electrophoresis. The washed slides were dehydrated with 70% ethanol, air-dried, and then kept at 4 ^°^C.

For agarose rehydration and DNA staining, 50 μl of PBS containing SYBR Staining Solution (Invitrogen) was added to the slides. The slides were analysed using an Olympus B × 50 fluorescence microscope at 400×. A minimum of 30 images were captured per sample. Comet Assay Software Project (CASP) software (version 1.2.3) was used to analyse the images.

### Artificial fertilization *in vitro*

To eliminate female impacts on paternal intergenerational inheritance, only one female was involved in the fertilization test. To ensure successful fertilization and meet the requirements for embryo sampling, 5 g of eggs (∼ 4000 eggs) were fertilized at a ratio of 500,000 spermatozoa per egg (Linhart et al. 2015) with 10 ml of activation solution (45 mM NaCl, 5 mM KCl, 30 mM Tris, pH 8.0, 160 mOsmol/kg) (Perchec et al. 1996). Prior to the fertilization experiment, spermatozoa concentration was measured using a Bürker cell hemocytometer (Marienfeld, Germany) with a final dilution of 5000. Fertilized eggs were cultured in Petri dishes at 21^°^C. The embryo samples at the mid-blastula stages were collected at 7.5 h post fertilization (HPF). Fertilization, hatching and malformation rates were calculated using the formula: fertilization rate (%) = fertilized eggs/total eggs *100, hatching rate (%) = hatched embryos/total eggs *100, and malformation ratio (%) = abnormal larvae/total hatched larvae *100.

### Sampling and DNA extraction

The DNA methylation level was assessed in fresh and stored sperm and in samples of mid-blastula embryos. In detail, a 1.8 ml cryotube containing 2 μl of each sperm sample was snapc:frozen in liquid nitrogen and kept at −80 c: until further use. The DNA isolation kit (Qiagen DNeasy, Hilden, Germany) was used to isolate sperm genomic DNA according to the manufacture’s protocol. The developed mid-blastula embryos (40-60 embryos/tube) from each treatment were sampled at 7.5 HPF. Before sampling, embryos were washed in triplicate with RNAse- and DNAse-free molecular grade water (Invitrogen). DNA from embryo samples was extracted using a modified SDS-based DNA extraction method according to (Natarajan et al., 2016). The DNA concentration was determined using a Nanophotometer Pearl (Implen, Munich, Germany). Finally, the gDNA’s purity and integrity were evaluated by electrophoretic separation and visual assessment in a 1.5% agarose gel.

### Assessment of global DNA methylation using isotope-dilution LC-MS/MS

Ten μg of DNA from each group of sperm and embryo samples were hydrolysed to 2’-deoxynucleosides using micrococcal nuclease from *Staphylococcus aureus*, bovine spleen phosphodiesterase and calf intestinal alkaline phosphatase (all from Sigma-Aldrich, Taufkirchen, Germany) as described previously (Schumacher et al. 2013). [^15^N_2_,^13^C_1_]-dC, and 5-mdC-d_3_ (both from Toronto Research Chemicals, Toronto, Canada) were used for quantification of dC and 5-mdC in DNA hydrolysates by isotope-dilution LC-MS/MS as reported recently (Gerecke et al. 2018). In brief, nucleoside analytes were separated using a Poroshell 120 EC-C8 column (3.0 x 150 mm, 2.7 µm) fitted in a 1290 Infinity II HPLC linked to an Ultivo (G6465) triple-quadruple mass spectrometer (all from Agilent Technologies, Waldbronn, Germany). Electrospray ionization (ESI+) was used to create protonated precursor ions, and the MS/MS detector was set to multiple reaction monitoring (MRM). For quantification, the following mass transitions were used (qualifier mass transitions are in parentheses): dC: m/z 228.1 → 112.0 (94.9); [^15^N_2_,^13^C_1_] dC: m/z 231.1 → 114.9 (97.9); 5-mdC: m/z 242.1 → 126.0 (109.1), and 5-mdC-d_3_: m/z 245.1 → 129.0 (112.0). Using the MassHunter program (*Quantitative Analysis* version 10.1) from Agilent Technologies, peak regions were assessed, and nucleoside concentrations in the DNA hydrolysates were computed based on the quantities of internal standards utilized.

### DNA library construction, whole-genome bisulfite sequencing, data quality control and mapping

Over 30 µg of total DNA were sent from each embryos sample (three males) to Admera Health Biopharma Services for WGBS analysis. In detail, isolated genomic DNA was quantified using the Qubit 2.0 DNA HS Assay (Thermo Fisher Scientific, Massachusetts, USA) and the quality was assessed by Tapestation genomic DNA Assay (Agilent Technologies, California, USA). Then, following the manufacturer’s protocol, bisulfite conversion was performed using the EZ DNA Methylation-Gold Kit (Zymo Research, California, USA), and DNA was fragmented to 300-400 bp. After the bisulfite conversion, library preparation was performed using Accel-NGS® Methyl-Seq DNA Library kit (Swift Biosciences, Michigan, USA). Thereafter, library yields were determined using the Qubit 2.0 DNA HS Assay and the quality of the libraries generated was assessed using the TapeStation HSD1000 ScreenTape (Agilent Technologies Inc., California, USA). The final libraries quantity was assessed by APA SYBR® FAST qPCR with QuantStudio® 5 System (Applied Biosystems, California, USA). Based on qPCR QC values, equimolar pooling of libraries was performed. A 20% PhiX Spike In was added to the samples to provide a bisulfite conversion efficiency and added during the sequencing to ensure sequencing quality. Finally, samples were sequenced on an Illumina® NovaSeq S4 (Illumina, California, USA) with a read length configuration of 150 PE for [188] M PE reads per sample (94M in each direction).

We conducted fundamental statistical analyses on the read quality of the raw data using FastQC (fastqc_v0.11.5). Thereafter, the read sequences generated by the Illumina pipeline in FASTQ format were then pre-processed using Trimmomatic software (Trimmomatic-0.36) with the parameter (SLIDINGWINDOW: 4:15; LEADING:3, TRAILING:3; ILLUMINACLIP: adapter.fa: 2: 30: 10; MINLEN:36). After filtering the reads, the remaining reads were considered as clean reads, and all subsequent analyses were conducted using this set of reads. Finally, upon completion of data cleaning, we utilized FastQC to generate basic statistical metrics that assessed the quality of the resulting reads.

Clean data were obtained after filtering and then the Bismark software (version 0.16.3; Krueger et al. 2011) was used to perform alignments of bisulfite-treated reads to a reference genome (assembly accession: GCF_000951615.1). The reference genome was firstly transformed into bisulfite-converted version (C-to-T and G-to-A converted) and then indexed using bowtie2 (Langmead and Salzberg 2012). Before being directly matched to similarly converted versions of the genome, sequence reads were also transformed into fully bisulfite-converted versions (C-to-T and G-to-A converted). The methylation status of every cytosine site in the sequence reads that result in a distinct best alignment from the two alignment procedures (original top and bottom strand) is then inferred by comparing them to the typical genomic sequence. The same reads aligned to the same regions of the genome were considered duplicated. Sequencing depth and coverage were summarised using deduplicated reads.

### Analysis of methylation levels and identification of DMRs

To identify the methylation site, we modelled the sum Mc of methylated counts as a binomial (Bin) random variable with the methylation rate. After the methylation sites were determined, the methylation level (ML) for each window or C site was calculated according to the formula ML = reads (mC)/reads (mC) + reads (umC). The motif of different sequencing contexts CpG, CHG and CHH (where H is A, C or T) was determined to reveal the sequence characteristics at the upstream and downstream regions of methylated cytosine (C) in the different context sequences. DMRs were identified using DSS software (parameter: smoothing = TRUE, smoothing.span = 200, delta = 0, p.threshold = 1e-05, minlen = 50, minCpG = 3, dis.merge = 100, and pct.sig = 0.5) (Park and Wu 2016). In detail, DMRs were classified as DMRs if they had a minimum length of 50 bp, contained at least three CpG sites, and were at least 50% statistically significant differentially methylated CpGs (*p* < 1e-05). The DMRs were ranked by the sum of the test statistics of the contained CpG sites. We determined the DMGs whose gene body region from transcriptional start sites (TSS) to transcription end sites (TES) or the promoter region (upstream 2 kb from the TSS) overlapped with the DMRs in accordance with the distribution of DMRs in the genome.

The term “differentially methylated genes” (DMGs) was used to describe whose promoter or gene body regions (upstream 2 kb from the transcription start sites, or TSS) overlapped the DMRs. Additionally, the methylation levels of genomic functional regions were also analysed. These regions included promoter, exon, intron, CGI (CpG island) and CGI shore (defined as the 2 kb sequence flanking a CGI). The average rate of methylation was calculated for each functional genomic region.

Gene Ontology (GO) enrichment analysis of genes associated with DMRs was performed using the GOseq R package (Young et al. 2010). GO terms with a corrected *P value* of less than 0.05 were considered significantly enriched to be DMR-related. We conducted Kyoto Encyclopedia of Genes and Genomes (KEGG) pathway enrichment analysis to identify significantly enriched metabolic or signalling pathways in genes with DMRs compared to the whole genome background. The statistical enrichment of DMR-related genes in the KEGG pathway was examined using the KOBAS program (Mao et al. 2005).

### Statistical analysis

The normality of the distribution and homogeneity of variances for all data were checked using Shapiro-Wilk and Levene’s tests. Statistical significance of sperm phenotypic parameters, mdC/dC (%) from LC-MS/MS was tested either using one-way ANOVA followed by a Tukey test or the nonparametric Kruskal–Wallis test followed by a Dunn pairwise comparison. All statistical analyses were performed with GraphPad Prism 9.1.0 (GraphPad Software, San Diego, CA, USA) and R programming language. Differences were considered significant at *p* < 0.05 except for special instructions.

## Results

### Short-term storage decreases sperm motility traits, DNA integrity and fertilizing ability

Comparison of biological traits of sperm between fresh and short-term stored sperm showed significant decreases in percentage of motility (Fig. 2A) and increase in percentage of sperm with DNA fragmentation (Fig. 2D) after 3 or 6 DPS (*p* < 0.05). However, a non-significant trend towards a decrease was observed for curvilinear velocity (VCL) (Fig. 2B), and a reduction in viability after 6 DPS (Fig. 2C). The fertilization and hatching rates were decreased when 6-day-stored sperm was used for fertilization (*p* < 0.05; Fig. 2F). Due to a high sperm: egg ratio used in the fertilization experiment, the fertilization and hatching rates were also high for short-term stored sperm (> 90%). Meanwhile, no changes in larval malformation were observed (*p* > 0.05; Fig. 2F). These results showed that storage-induced decreases in sperm quality was accompanied by fertility loss, although it did not have an apparent effect on embryonic development to hatching.

**Fig. 2.**
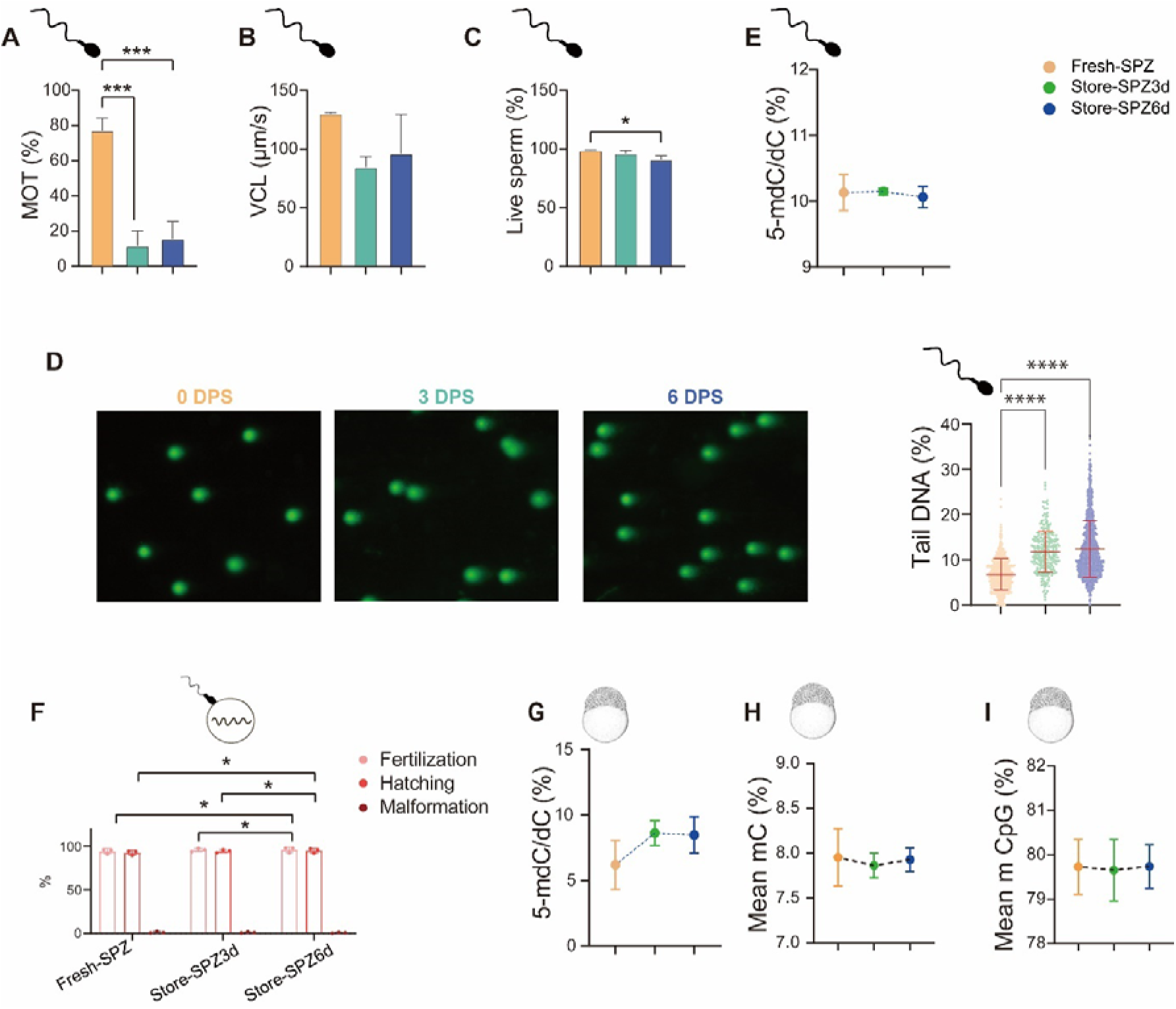
Effects of short-term storage on sperm quality, fertility and global DNA methylation. (**A**) percent of spermatozoa motility; (**B**) curvilinear velocity (VCL) of motile spermatozoa; (**C**) percentage of live spermatozoa with intact membrane; (**D**) DNA fragmentation levels (defined by the percent of tail DNA); (E) fertilization and hatching rates of short-term stored sperm; (F) malformation rate in the embryos; (**G), (H**) global DNA methylation [5-mdC/dC (%)] of sperm and embryos at mid-blastula; (**I**) average methylation level of all the genome cytosine sites of mid-blastula; (**J**) the average methylation level of all the genome CpG-context cytosine sites of mid-blastula. Statistical significance was determined using one-way ANOVA followed by a Tukey test; data of sperm, embryos and fertilization from each group (control group: Fresh_SPZ; 3-day-sperm storage group: Store_SPZ3d and 6-day-sperm storage group: Store_SPZ6d) represent the mean ± S.D. of three biological replicates (* *p* < 0.05; *** *p* < 0.001; **** *p* < 0.0001).

### Short-term storage does not alter global DNA methylation in sperm and embryos

The 5-mdC/dC in short-term stored sperm and their embryos remained unchanged (*p* > 0.05; Fig. 2E, G). Also, mean mC and mCpG in the embryos derived from short-term stored sperm were similar to those derived from fresh sperm (*p* > 0.05; Fig. 2H, I). Taken together, these results indicated that short-term storage decreased sperm quality, but had no significant effects on global DNA methylation of the derived embryos.

### Data generation and mapping to the reference genome

To reveal the differences in the DNA methylation at the whole genome level of the mid-blastula embryos derived from fresh and short-term stored sperm, we performed WGBS of embryos with a mean yield of 188 million reads. After quality control, 48–58% of the reads were uniquely mapped to the reference genome, and an average of 90% base covered ≥ 5×. The coverage depths in the embryos from fresh and stored sperm were at least above 7.5 (Supplementary Table S1).

### DNA methylation in the sequencing contexts

The specific DNA methylation level of the three sequence contexts, CpG, CHG and CHH, was analysed for the embryos derived from fresh and stored sperm samples. The average methylation levels at C-sites in the embryos originating from short-term stored sperm were similar; 7.95%, 7.86% and 7.93% for fresh (Fresh_SPZ), 3-day-stored sperm (Store_SPZ3d) and 6-day-stored sperm (Store_SPZ6d), respectively (Fig. 2H and Table 1). The mean methylation levels of CpG were higher than CHG, and CHH in the embryos that originated from fresh or short-term stored sperm. The values were 79.73%, 0.93% and 1.01% in the embryos obtained from fresh sperm, 79.65%, 0.78% and 0.86% in the embryos obtained from 3-day-stored sperm, 79.74%, and 0.90% and 1.00% in the embryos obtained from 6-day-stored sperm (Table 1). Since most mC sites covered by CpG contexts, only DMRs in CpG contexts were considered for subsequent analyses. The methylation level in the CpG context did not differ among the embryos obtained from short-term stored sperm (*p* > 0.05; Fig. 2I). The CpG methylation on each chromosome in the embryos from stored sperm varied from that in the embryos originating from the fresh sperm (Supplementary Fig. S1).

**Table 1.**
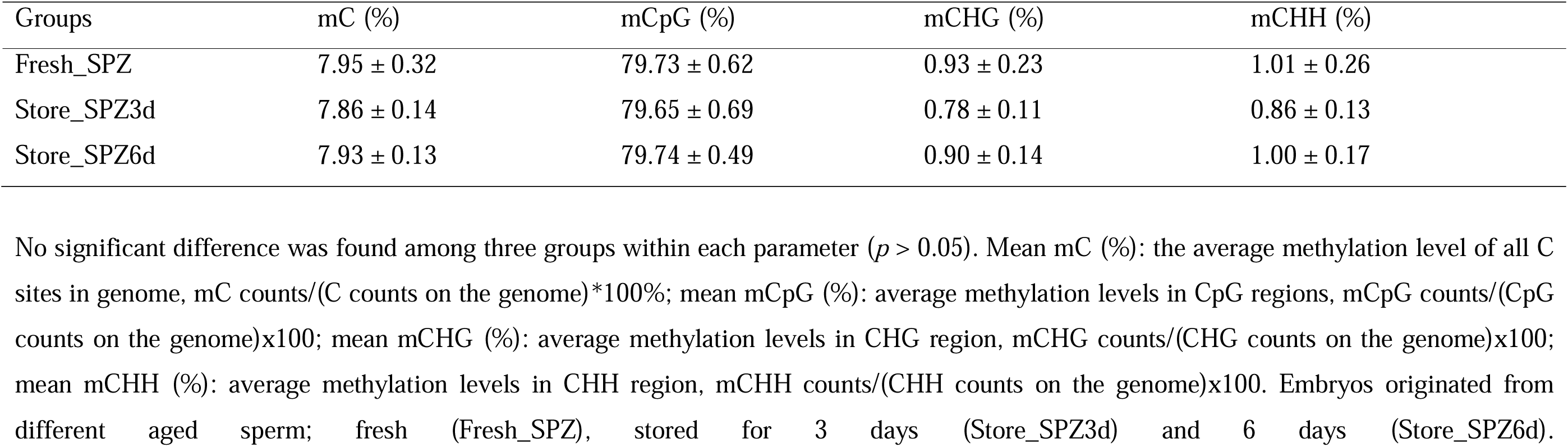
Percentage of total genome methylation statistics. Data represent means ± S.D. (*n* = 3 fish replicates)

| Groups | mC (%) | mCpG (%) | mCHG (%) | mCHH (%) |
| --- | --- | --- | --- | --- |
| Fresh_SPZ | 7.95 ± 0.32 | 79.73 ± 0.62 | 0.93 ± 0.23 | 1.01 ± 0.26 |
| Store_SPZ3d | 7.86 ± 0.14 | 79.65 ± 0.69 | 0.78 ± 0.11 | 0.86 ± 0.13 |
| Store_SPZ6d | 7.93 ± 0.13 | 79.74 ± 0.49 | 0.90 ± 0.14 | 1.00 ± 0.17 |

No significant difference was found among three groups within each parameter ( $p > 0.05$ ). Mean mC (%): the average methylation level of all C sites in genome, mC counts/(C counts on the genome)\*100%; mean mCpG (%): average methylation levels in CpG regions, mCpG counts/(CpG counts on the genome)x100; mean mCHG (%): average methylation levels in CHG region, mCHG counts/(CHG counts on the genome)x100; mean mCHH (%): average methylation levels in CHH region, mCHH counts/(CHH counts on the genome)x100. Embryos originated from different aged sperm; fresh (Fresh\_SPZ), stored for 3 days (Store\_SPZ3d) and 6 days (Store\_SPZ6d).

According to cluster analysis based on CpG methylation levels, the samples were distributed across three clusters, each containing two, three and four samples. However, the samples of various stored sperm groups did not appear to cluster together (Fig. 3A). Meanwhile, a heat map of DNA methylation based on the same criteria showed a very high correlation between the treatments within the same male (Pearson’s correlation coefficient > 0.94) (Fig. 3B).

**Fig. 3.**
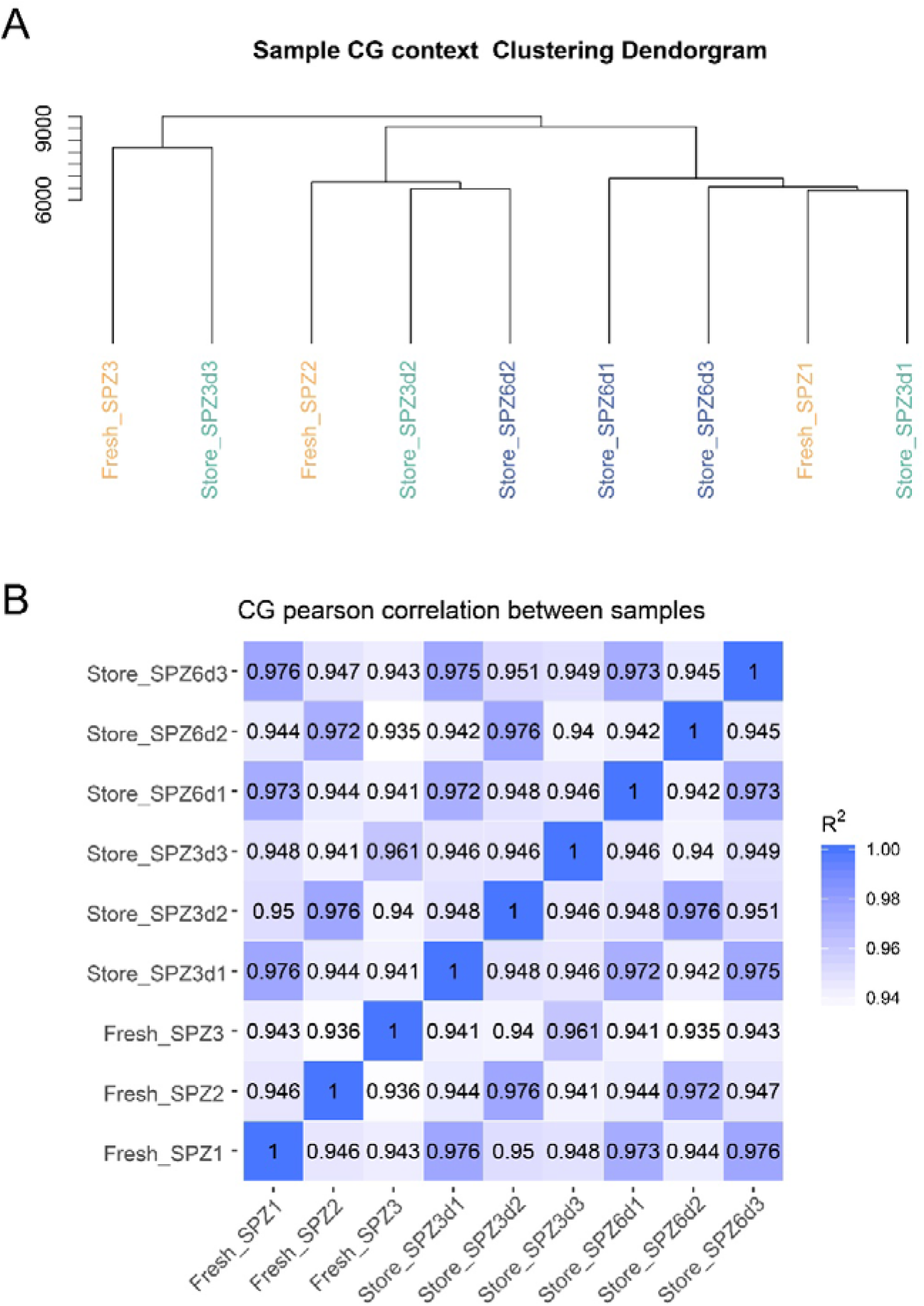
Clustering and correlation of analysis of samples based on CpG methylation level. (**A**) Hierarchical clustering by CpG methylation levels in common carp embryos obtained from fresh and short-term stored sperm; (**B**) heat map and correlation analysis based on CpG data among common carp embryos obtained from fresh and short-term stored sperm. Numbers in each cell represent the pairwise Pearson’s correlation scores. Data obtained from sperm of three males with different storage time: fresh, 3-days- and 6-days-stored sperm from male 1 (Fresh_SPZ1, Stored-_SPZ3d1, Stored-_SPZ6d1), male 2 (Fresh_SPZ2, Stored-_SPZ3d2, Stored-_SPZ6d2), and male 3 (Fresh_SPZ3, Stored-_SPZ3d3, Stored-_SPZ6d3).

### DNA methylation in the genomic elements

DNA methylation levels in different genomic elements were analysed in the embryos produced from fresh and short-term stored sperm (Fig. 4). Generally, the average CpG methylation levels were high in the CpG island and CGI shore and low in the 5’ untranslated region (5’ UTR) and promoter regions (Fig. 4A-C). Comparison between embryos showed that (a) the average methylation levels in several genomic regions (such as CpG island, CGI shore, promoter and 5’ UTR regions) were slightly higher in the embryos from fresh sperm than those of 3-day-stored sperm (Fig. 4A), (b) the average methylation levels in genomic regions of CpG island were higher (although not in the promoter region) in the embryos of 3-day-stored sperm than those of 6-day-stored sperm (Fig. 4B), and (c) the average methylation levels showed similar trends in the embryos of fresh sperm *vs.* those of 3-day-stored sperm (Fig. 4C). Analyses of DNA methylation levels in the genome upstream, gene body and downstream showed lower CpG methylation at upstream and downstream regions compared to that in the gene body region (Fig. 4D-F). In this context, the CpG methylation levels in all these regions in the fresh sperm were similar as those in the 3-day-stored sperm (Fig. 4D). Also, the CpG methylation levels of upstream and downstream regions in the embryos of fresh sperm and 3-day-stored sperm were lower than embryos of 6-days-stored sperm (Fig. 4E, F).

**Fig. 4.**
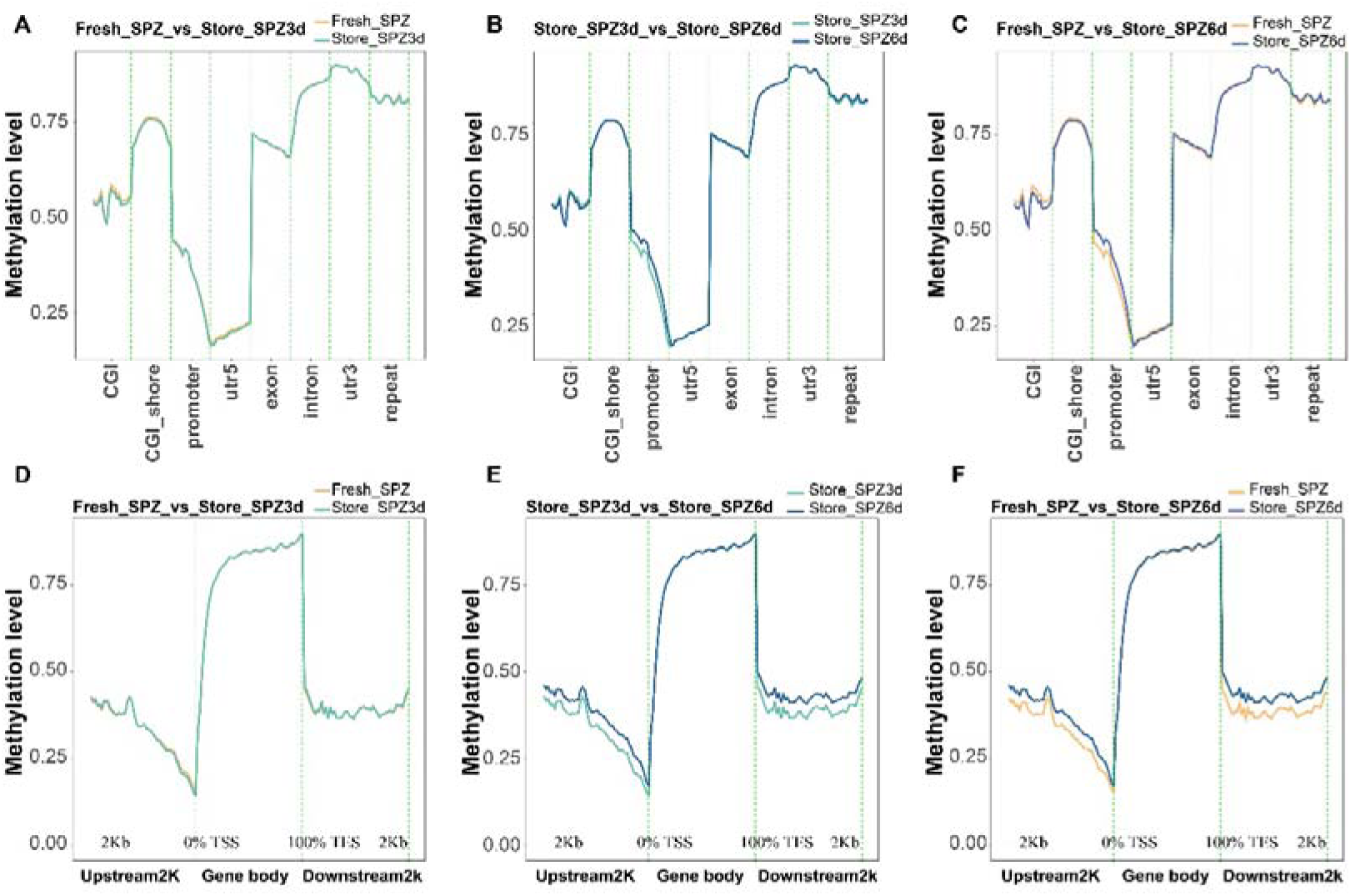
Comparison of methylation levels on different genomic elements and distribution of up-stream and down-stream among embryos derived from fresh sperm (Fresh_SPZ), stored sperm for 3 days (Store_SPZ3d) and for 6 days (Store_SPZ6d) *in vitro*. (**A)-(C**) Density plot of methylation levels on different genomic elements in CpG; (**D)-(F**) density plot of methylation levels on different regions of up-stream, gene body and down-stream. (*n* = 3 for each group).

### DMRs among embryos developed from fresh and stored sperm

In total, 4761 DMRs with annotation were identified between the embryos of fresh and embryos of 3-days-stored sperm, including 2734 hyper-DMRs in fresh sperm and 2027 hypo-DMRs. There were 8624 between embryos of 3-day-stored sperm and embryos of 6-day-stored sperm (4069 hyper-DMRs and 4555 hypo-DMRs) and 7726 DMRs between embryos of fresh and 6-day-stored sperm (4062 hyper-DMRs and 3664 hypo-DMRs) (Supplementary Fig. S2). The average DMR methylation level and distribution pattern were similar between embryos of fresh sperm and 3-day-stored sperm (Fig. 5A). However, there were more hyper- and hypo methylation in the embryos of 6-day-stored sperm than those obtained from fresh and 3-day-stored sperm (Fig. 5B, C).

**Fig. 5.**
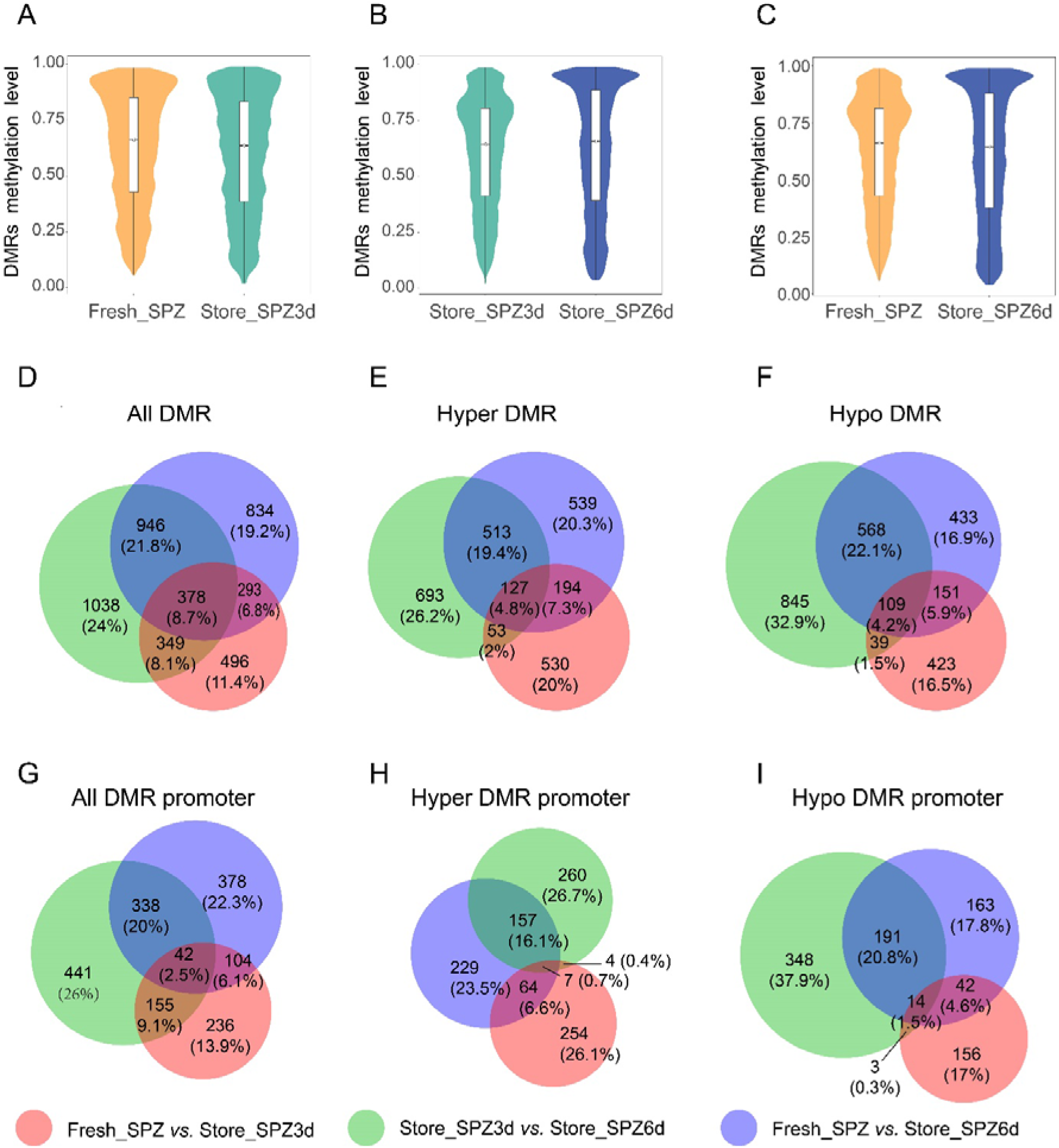
Analysis of differential methylation levels and regions (DMRs). (**A)-(C**) Violin plots of methylation levels of DMRs in the embryos original from fresh sperm (Fresh_SPZ), 3-day-stored sperm (Store_SPZ3d) and 6-day-stored (Store_SPZ6d) *in vitro*. All comparisons: *P* < 0.0001. (**D)-(F**) Venn diagrams of DMR-anchored genes in gene region including all types, hyper DMR and hypo DMR-target genes in CpG content of embryos originating from fresh sperm (Fresh_SPZ), 3-day-stored sperm (Store_SPZ3d) and 6-day-stored sperm (Store_SPZ6d) *in vitro*. (**G)-(I**) Venn diagrams of DMR promoter-target genes including all types, hyper DMR and hypo DMR-target genes in CpG content of embryos original from fresh sperm (Fresh_SPZ), 3-day-stored sperm (Store_SPZ3d) and 6-day-stored sperm (Store_SPZ6d).

In the embryos of fresh sperm group *vs.* 3-day-stored and 6-day-stored sperm at all different gene regions (CGI, CGI shore, promoter, TSS, 5’ UTR, exon, intron, 3’ UTR and TES), the number of hyper-methylated DMRs was also significantly higher than hypo-methylated DMRs (Supplementary Fig. S3A, C). However, an opposite trend was observed between 3-day- and 6-day-stored sperm (Supplementary Fig. S3B). By anchoring the DMRs to the gene body (from TSS to TES) and promoter region, 2053, 3687 and 3313 differentially methylated genes (DMGs) in embryos from fresh *vs.* 3-days-stored sperm, 3-days-stored sperm *vs.* 6-days-stored sperm, and fresh *vs*. 6-days-stored sperm, respectively, were identified from CpG-type DMRs (Fig. 5D-I).

A total of 1516, 2711 and 2451 differentially methylated genes (DMGs) in the gene body was identified between embryos from fresh sperm *vs*. 3-day-stored sperm, 3-day-stored sperm *vs*. 6-day-stored sperm, and fresh sperm *vs*. 6-day-stored sperm (Fig. 5D), including 904, 1386 and 1373 upregulated DMGs (Fig. 5E) and 722, 1561 and 1261 downregulated DMGs, respectively (Fig. 5F). Moreover, 321 and 206 common DMGs were either upregulated or downregulated in the embryos of fresh sperm compared to those of 3-day-stored and 6-day-stored sperm (Fig. 5E, F). In the promoter regions, more common DMGs (56) were identified in downregulated genes than upregulated genes (11) in the above comparisons (Fig. 5G-I).

### Functional enrichment analysis for genes with differentially methylated genes (DMGs)

To reveal the biological functions of differentially methylated genes among embryos derived from fresh and stored sperm, we performed pathway analysis of differentially methylated genes based on the GO and KEGG database. Five GO terms were notably enriched from GO enrichment analysis, and the results indicated that promoter hypo-DMGs in the embryos only from sperm groups of fresh *vs*. 6-days-stored were enriched in the biological process related to cell-cell adhesion and homophilic cell adhesion (Table 2, Supplementary Table S2). However, in the analysis of promoter hyper-DMGs, hyper- and hypo-DMGs in the gene body, there was no significant differences in the genes of the biological process (BP), cellular component (CC) and molecular function (MF) classifications among three groups (*p* >0.05; Supplementary Fig. S4 and S5).

**Table 2.**
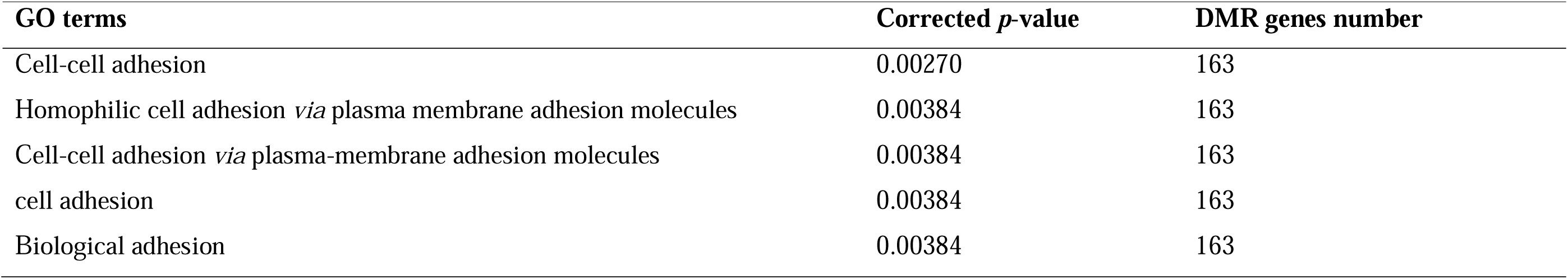
GO enrichment analysis of promoter hypo-DMGs (top five terms) in embryos from fresh sperm *vs*. 6-day-stored sperm.

In addition, KEGG enrichment analysis showed only oxidative phosphorylation was significantly enriched in promoter hyper-DMGs between fresh and 3-day-stored sperm groups, and hypo-DMGs when compared with 3-day-stored and 6-day-stored sperm groups (Supplementary Table S3). The gene body hyper-DMGs were enriched in KEGG pathways, such as neomycin, kanamycin and gentamicin biosynthesis, calcium signalling pathway, adrenergic signalling in cardiomyocytes, regulation of actin cytoskeleton and melanogenesis between fresh and 6-day-stored sperm (Fig. 6C) and pathways related to focal adhesion, cell adhesion molecules (CAMs) and RNA transport enriched in gene body hyper-DMGs between 3-day-stored and 6-day-stored sperm (Fig. 6B). In the gene body hypo-DMGs, pathways included but were not limited to starch and sucrose metabolism, neomycin, kanamycin and gentamicin biosynthesis, galactose metabolism, Apelin signalling pathway and RNA transport (fresh sperm *vs*. 3-day-stored sperm; Fig. 6D); MAPK signalling pathway and Adherents junction (fresh sperm *vs.* 6-day-stored sperm; Fig. 6E). Only three pathways of cardiac muscle contraction, cell adhesion molecules (CAMs) and adrenergic signalling in cardiomyocytes were different between 3-day-stored sperm and 6-day-stored sperm *in vitro* (Fig. 6F). Detailed information on the KEGG enrichment analysis is shown in Supplementary Table S3.

**Fig. 6.**
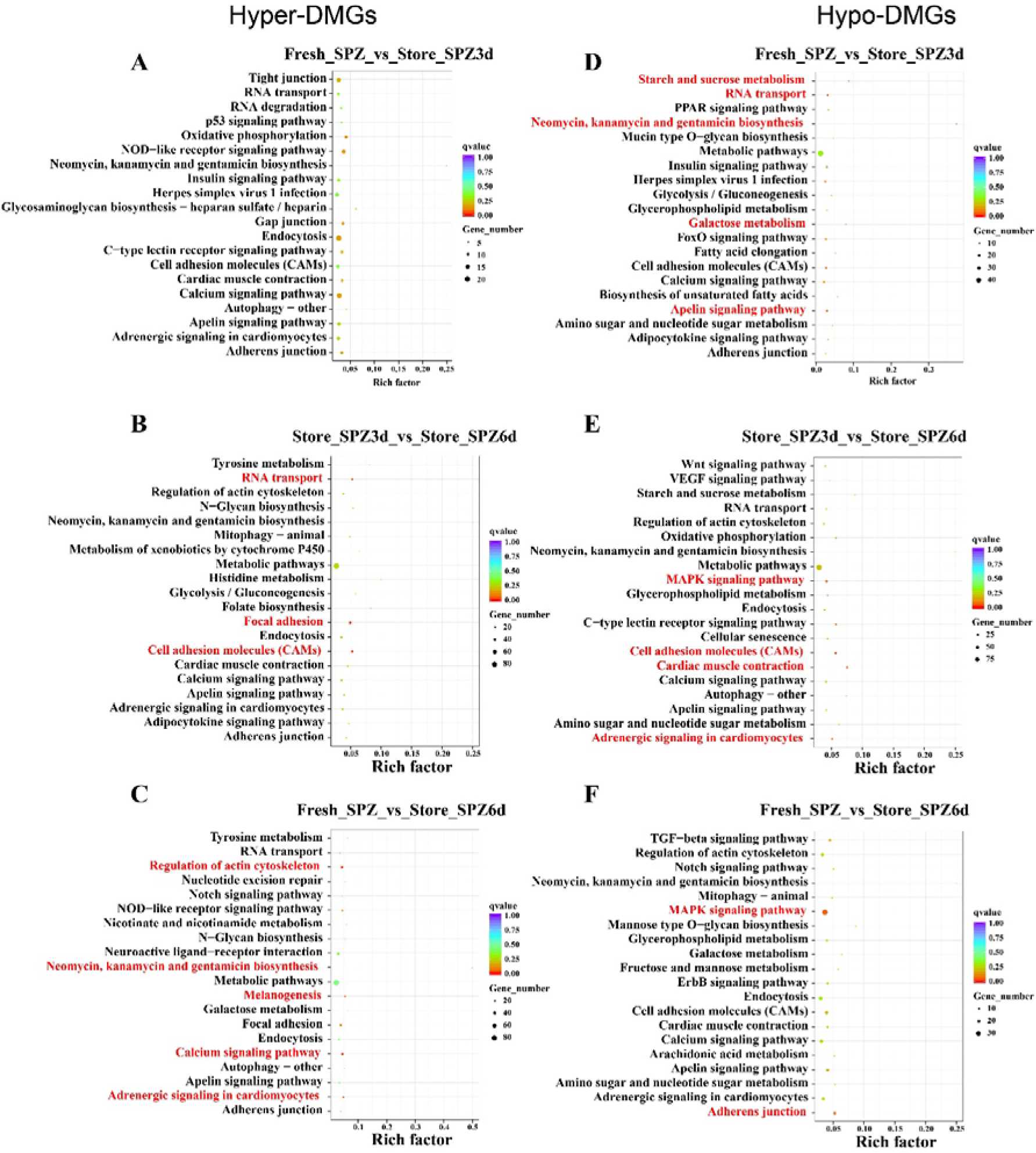
(**A)-(C**) KEGG enrichment analyses of DMGs in gene body hyper-DMGs and (**D)-(F**) hypo-DMGs related pathways with the most significant *P* value of different methylation genes among embryos derived from fresh sperm (Fresh_SPZ), 3-day-stored sperm (Store_SPZ3d) and 6-day-stored sperm (Store_SPZ6d) *in vitro*. The *x*-axis indicates percentages of differential methylated genes belonging to the corresponding pathway. The sizes of bubble represent the number of DMGs in the corresponding pathway, and the colours of the bubble represent the enrichment *P*-value of the corresponding pathway. Red colour shows the significant pathways with corrected *p*-value < 0.05.

## Discussion

DNA methylation is one of the most important epigenetic modifications involved in regulating growth, reproduction, disease resistance and stress response in aquaculture species (Liu et al. 2022). However, little is known about its function when short-term stored sperm is used for artificial fertilization in hatcheries. Fish sperm, like those of other vertebrates (Darszon et al. 1999), are immobile in the sperm duct and spermatozoa can be stored in the natural or artificial seminal plasma in a quiescent state for a short time without loss of motility and fertilizing capacity (Shazada et al. 2023). In the present study, the main objective was to investigate DNA methylation profiles of embryos produced from short-term stored common carp sperm using WGBS. We also assessed sperm functions after the short-term storage. Taken together, sperm motility characteristics, DNA integrity and viability were reduced, but global DNA methylation of spermatozoa and resulting embryos were not affected.

Sperm motility is an essential phenotype to manifest fertilization capacity and pathway to offspring formation. In this study, the method of sperm storage was similar to that used in our previous studies (Cheng et al. 2022a, b; Zhang et al. 2023a). Sperm motility traits were measured immediately after storage, and results were consistent with our recent study (Zhang et al. 2023a), which indicated that the percentages of motile spermatozoa and spermatozoa viability were significantly decreased after storage at 6 DPS even with elevated temperature after sperm storage. No significant decrease in spermatozoa velocity was observed after 6 DPS compared to fresh sperm, which is consistent with that of the fertilization rate. This is due to the fertilization rate being primarily dependent on sperm velocity (Linhart et al. 2005). However, the storage procedure must be reliable in order to preserve the genetic and epigenetic contents of sperm during the storage period. In terms of sperm DNA integrity, there was an increased level of DNA fragmentation in 6-day-stored sperm with respect to the tail DNA (Fig. 2). This suggests that short-term storage causes damage to sperm DNA in common carp, which is consistent with previous studies in mammals and fishes (Pérez-Cerezales et al. 2009; Shaliutina et al. 2013; Khezri et al. 2019). However, the stability of sperm chromatin shows species specificity during *in vitro* storage, for example, DNA damage was increased in Russian sturgeon (*Acipenser gueldenstaedtii*), Siberian sturgeon (*Acipenser baerii*) and rainbow trout (*Oncorhynchus mykiss*) sperm at 6-, 3- and 2-days post-storage, respectively (Pérez-Cerezales et al. 2009; Shaliutina et al. 2013). DNA fragmentation and sperm membrane integrity have been reported to be associated with oxidative stress (Gazo et al. 2015). In addition, in terms of global DNA methylation, the current result is consistent with a previous study, which suggested that sperm preservation in extenders can, to a large extent, mitigate the adverse effects of ROS and safeguard the cell structure.

In general, the contexts of DNA methylation have been identified as CpG, CHH and CHG, but in animal genomes, DNA methylation mostly occurs on the cytosines of a CpG dinucleotide (Lee et al. 2010). In fishes, high levels of CpG methylation have been reported, such as 94% in zebrafish (Potok et al. 2013), 98% in grass carp (*Ctenopharyngodon idellus*) (He et al. 2022), 71-73% in Chinese perch (*Siniperca chuatsi*) (Pan et al. 2021), 98% in tiger pufferfish (*Takifugu rubripes*) (Zhou et al. 2019), and 76-78% in ricefield eel (*Monopterus albus*) (Chen et al. 2021). In this study, the average global methylation level in a CpG context across the nine different common carp embryo samples was 79.71% (Table 2), and a lower methylation level was observed in the CHH (0.86-1.01) and CHG (0.78-0.93). It was similar to our previous study which showed 79% mCpG in common carp spermatozoa (Cheng et al. 2021). Generally, DNA methylation levels vary depending on the specific tissue, sex and stage of development (Strömqvist et al. 2010; Wang and Bhandari 2019). Based on the changes in sperm DNA fragmentation and from a previous study in sperm DNA methylome (Cheng et al. 2021), the question is raised of whether short-term induced damage to sperm DNA affects offspring development. Clustering analysis and Pearson correlation based on CpG methylation levels showed no distinct clustering and a very high positive correlation between males (Fig. 3). These findings show slight variation between the treated samples and imply that sperm storage-induced-specific differences in methylation of common carp embryos are small. Since the embryo samples from the different stored sperm belonging to the same male displayed similar methylation levels, genetic diversities may have a greater effect on DNA methylation than gamete manipulation.

Analysis of DMRs revealed a considerable increase in the number of hyper/hypo DMRs and DMRs target genes in the Fresh_SPZ *vs*. Store_SPZ6d and Store_SPZ3d *vs*. Store_SPZ6d comparison compared to the Fresh_SPZ *vs*. Store_SPZ3d comparison (Fig. 5). The association between sperm manipulation and sperm DNA methylation is inconsistent in the literature. Most of the studies have investigated long-term (cryopreserved) stored sperm. For example, cryopreservation has been shown to have effects on epigenetic markers that vary among animals (Chatterjee et al. 2017), cryopreservation agents (Depincé et al. 2020), and accuracy of the methods used in the investigation (Kurdyukov and Bullock 2016). Some studies have also shown that the influence of short-term storage of sperm on DNA methylation is due to the different levels of DNA fragmentation, and there is an increase in the number of downstream genes and their associated regulatory pathways that may be impacted with increasing DNA fragmentation in spermatozoa (Khezri et al. 2019). Sperm DNA fragmentation significantly correlates with field fertility performance and abnormal embryo development (Gosálvez et al. 2014). On the other hand, the sperm methylome is already determined during spermatogenesis; but there are factors that can mutate the DNA methylome of sperm, including the DMRs target genes. For example, there is evidence showing that oxidative stress produced during storage leads to changes in DNA methylation patterns in spermatozoa (Khezri et al. 2019). Oxidative stress not only leads to DNA fragmentation but also causes damage to sperm membranes and mitochondria, resulting in sperm with abnormal morphology and flagella. Consequently, sperm motility and the number of normal spermatozoa decreases. The higher frequency of DNA fragmentation observed in spermatozoa after 3 and 6 days of storage in this study may be due to possible elevated levels of oxidative stress (Fig. 2E).

The regional analysis showed that most of the DMRs were annotated to be present in the intron, CpG islands (CGI) and CGI shore regions (Fig. S3). Most organisms have intron sections, which can be present in both protein-coding genes and non-coding genes (Boivin et al. 2018). Methylation in gene promoters has drawn the greatest attention in the conventional model because it is frequently linked to transcriptional silence. However, recent research indicates that in some specific genes, the first intron’s methylation is related to gene expression (Anastasiadi et al. 2018). In accordance with the present results, the methylation pattern was similar to previous studies on Chinese perch, which showed the most DMR was concentrated in the intronic regions (Pan et al. 2021). Meanwhile, intronic enhancers that interact with the promoters of the relevant genes could partially explain it. Additionally, CGIs are short-interspersed DNA sequences with a higher CpG frequency than other regions in the genome. At the same time, CGI shores (up to 2 kb distant) are the immediate regions flanking CGIs (Deaton and Bird 2011). CGI methylation is frequently linked to transcriptional silence because CGIs are typically found near transcription start sites (Jones 2012). In the present study, embryos obtained from fresh sperm had slightly higher methylation levels than those obtained from stored sperm (Fig. 3), suggesting that stored sperm induce some genes transcription activity during embryogenesis.

By overlapping the differentially methylated regions and functional gene regions, 378 differentially methylated genes were identified among three embryo groups in this study. In addition, 1373 significantly hypo-DMGs were identified in the methylome analysis. GO enrichment analysis identified the biological process of differentially methylated genes between the embryos produced by fresh and stored sperm. These DMGs were involved in GO terms of cell adhesion. The cell adhesion molecules (CAMs) pathway was also enriched in KEGG analysis. Cell-cell adhesion molecules play crucial roles, especially in the early stages of reproduction, such as gamete transport, sperm-oocyte interaction, embryonic development and implantation (D’Occhio et al. 2020). The results from the current study suggest that prolongation of storage time may have higher impact on embryonic development. In rainbow trout embryos, genes belonging to cell adhesion are important in vascular development (Vehniäinen et al. 2016). Notably, studies have demonstrated that fertilization with spermatozoa exhibiting oxidative stress is associated with aberrant early cleavage events and changes in embryo transcript abundance. Mammalian embryos produced from ROS-treated sperm were identified with a similar pathway (cell adhesion) associated with differentially expressed genes (Burruel et al. 2013). Sperm preservation in extenders may also directly affect the epigenome of the paternal DNA. For instance, a recent study has shown that boar (*Sus scrofa*) sperm after preservation display various increases in the DNA fragmentation index (DFI) among individuals, and genes corresponding to low-high DFI comparison were associated with crucial processes such as membrane function, metabolic cascade and antioxidant defence system (Khezri et al. 2019). In the present study, it is possible that 6- days-stored sperm underwent oxidative stress according to an increase in the DNA fragmentation level (Fig. 2E). This may be the reason behind the observed induced changes in the epigenome of embryos resulting from 6-days-stored sperm.

Studies associated with the DNA methylome and transcriptome of the sperm following short-term storage and their sired offspring are few compared to cryopreserved sperm. The transcriptome and methylome analyses of cryopreserved sperm of viviparous black rockfish (*Sebastes schlegelii*) revealed that the cryopreservation affected the expression of 179 genes and methylation of 1266 genes in spermatozoa. These genes associated with differential expression or methylation were highly enriched in KEGG pathways of phospholipase D signalling pathways and xenobiotic and carbohydrate metabolism pathways, and GO terms of death, G-protein coupled receptor signalling pathways, and response to external stimulus (Niu et al. 2022). Fertilization with frozen-thawed sperm in mammals downregulates genes relevant to early embryo development, such as in biological pathways related to oxidative phosphorylation, DNA binding, DNA replication and immune response (Ortiz-Rodriguez et al. 2018). In the case of short-term stored sperm, it has been observed that transcripts of genes involved in energy metabolism and stress response were decreased gradually (Yang et al. 2021). These findings suggest that disruption to the transcriptome and DNA methylome of preserved sperm may have an adverse impact on the development of embryos.

In our previous study, more than thousands of DMRs (875 methylated and 854 unmethylated) were found between fresh and 96 h-stored common carp sperm (Cheng et al. 2021). In the present study, under the protection function of extenders, various KEGG pathways were identified from embryos obtained with 3-day-stored (72 h) and 6-day-stored (144 h) sperm, and higher changes in genes related to DMRs were identified following 6 days of storage than 3 days of storage. For example, at 6 days of post storage, KEGG pathways suggested short-term storage influenced calcium signalling, adrenergic signalling in cardiomyocytes, regulation of actin cytoskeleton, and melanogenesis in the resulting embryos. However, 3 days of sperm storage changed the pathways involved in metabolism, Apelin signalling and RNA transport, which play an irreplaceable role in the regulation of the heart, lymphatic development and precise spatiotemporal control of gene expression. Taken together, these results suggest that short-term storage of sperm may affect embryonic development in these aspects. For example, Fang et al. (2016) showed that DNA methylation played a crucial role in embryonic cardiomyocytes because the knockdown of DNA methyltransferase 3a was found to alter gene expression and inhibit the function of embryonic cardiomyocytes. Additionally, the crucial and varied roles performed by actin regulation during morphogenesis, including signals regulating by the Rho Family of Small GTPases and MAP Kinase Cascades have been found (Woolner and Martin 2006). Similar pathways in sperm after treatment with ROS have previously indicated changes in transcript abundance for genes related to actin cytoskeleton organization compared to good sperm (Burruel et al. 2013). In addition, melanogenesis is the process by which melanin, a pigment responsible for coloration, is produced and deposited in various cells and tissues. The functions of melanogenesis in fish embryos may involve camouflage, UV protection, temperature regulation, immune defence, oxygen balance and cell signalling and development. Sperm itself cannot directly regulate melanin production in embryos, but sperm provides paternal genetic information, including genes related to melanin production. These genes can be expressed when an embryo is developing, and they take a role in controlling the routes and procedures involved in the production of melanin. However, these results are insufficient to indicate the influence of stored sperm on the resulting embryos. Multiple factors from sperm, such as histone modifications and noncoding RNA, determine the gene expression and protein synthesis in embryos. Thus, the systemic examination of the epigenetic status and other multi-omics simultaneously is warranted.

## Conclusion

The current work provides evidence that sperm short-term storage may impact DNA methylation alterations in the resulting embryos. Prolongation of storage duration shows higher impacts on the DNA methylation of the derived embryos. Furthermore, DMR analysis revealed that genes involved in cell adhesion, calcium signalling, mitogen-activated protein kinase (MAPK) and adrenergic signalling, metabolism, and RNA transport were primarily affected in embryos derived from sperm stored for 6 days *in vitro*.

This is the first investigation elaborating on the single-base DNA methylome as an epigenetic alteration in embryos originating from the short-term stored sperm samples in common carp. Using common carp as a fish model organism provided us with an advantage to perform multi-stripping of a male and to collect enough volume of sperm to analyse sperm motility, DNA integrity, viability, epigenetics and to conduct fertility tests together. The current study reports enough robust data from the 180 mid-blastula stage embryos in each group of fresh sperm, and sperm stored for 3 and 6 days *in vitro* obtained from the same male’s sperm samples under either fresh or stored conditions. Our results provide valuable information on genetic management of broodstock of common carp in aquaculture. However, the parallel transcriptomic analysis in future research would be informative and may reveal further consequential epigenetic changes in the resulting progeny from the short- or long-term stored gametes.

## Abbreviations

LC-MS/MS: liquid chromatography with tandem mass pectrometry
WGBS: Whole-genome bisulfite sequencing
DPS: days post stripping
DMRs: differentially methylated regions
DMG: differentially methylated genes
GO: gene ontology
KEGG: Kyoto Encyclopaedia of Genes and Genomes
Fresh_SPZ: fresh sperm
Store_SPZ3d: *in vitro* stored sperm for 3 days
Store_SPZ6d: *in vitro* stored sperm for 6 days
TSS: transcriptional start sites
TES: transcription end sites
5’ UTR: 5’ untranslated region.

## Acknowledgements

The authors thank Susanne Scheu (Freie Universität Berlin) for her excellent technical assistance with the DNA work-up for LC-MS/MS measurements.

## Funding

This study was funded by the Ministry of Education, Youth and Sports of the Czech Republic (LRI CENAKVA, LM2023038), by project Biodiversity (CZ.02.1.01./0.0/0.0/16_025/0007370), by the Grant Agency of the University of South Bohemia in České Budějovice (107/2022/Z), by the Czech Science Foundation (23-06426S) and by the National Agency for Agriculture Research, Czech Republic (QK21010141).

## Competing interests

The authors declare that they have no competing interests.

## Availability of data and materials

Data will be made available on request.

## Authors’ contributions

YC and OL conceived and designed the experiments. OL supervised the study and provided the funding. YC, SZ, PV, FS, IG, BK, AD, VR, MR and ZL carried out the experiments and collected data. YC and RN performed statistical analyses and generated figures and tables. YC, RN, SGW, SMHA and OL wrote the manuscript. YC, SZ, RN, PV, PL, JS, SMHA, CL and OL revised and reviewed the manuscript. All authors read and approved the final version of the manuscript.

## Supplementary Files

**Table S1.**
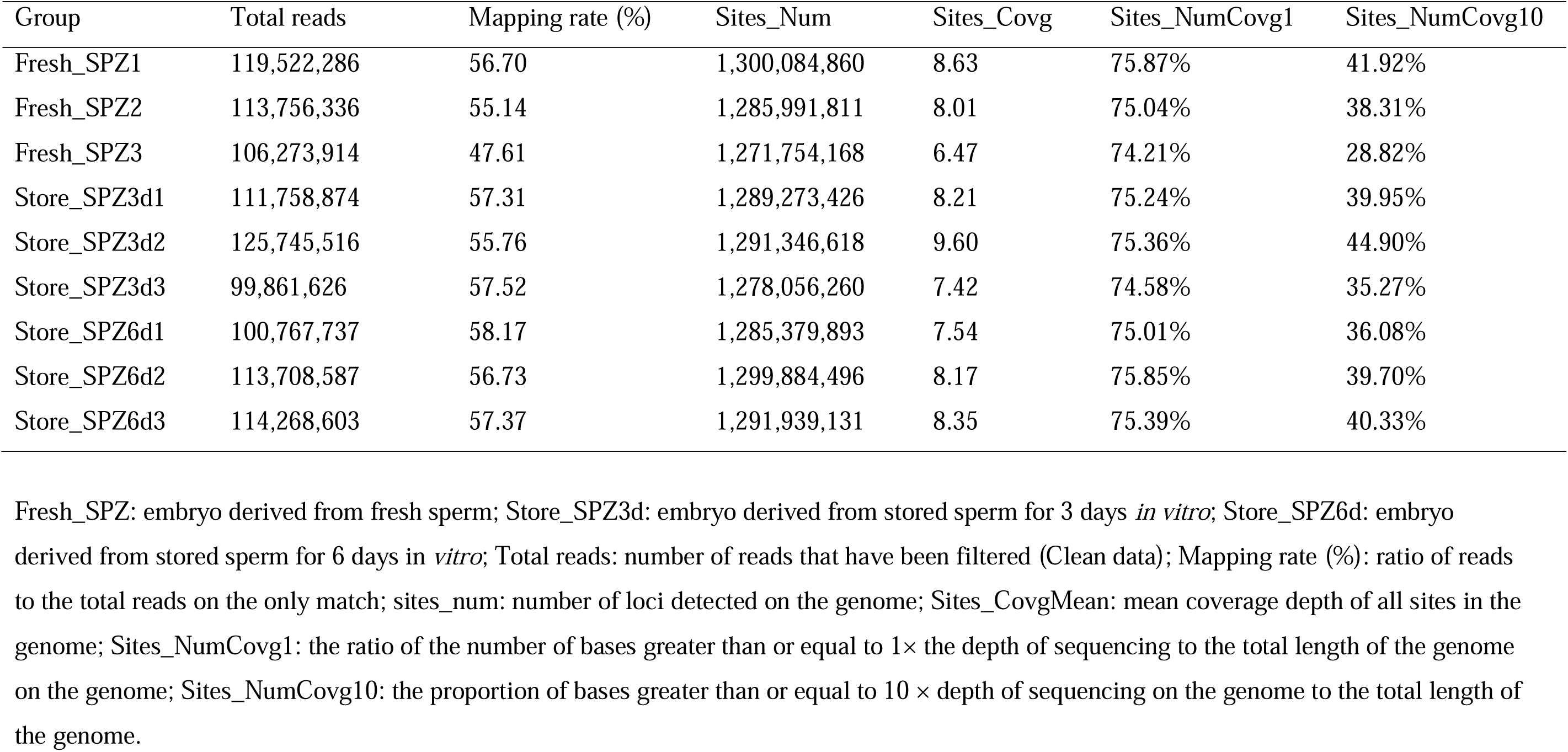
Genome coverage statistical analysis.

**Table S2.**
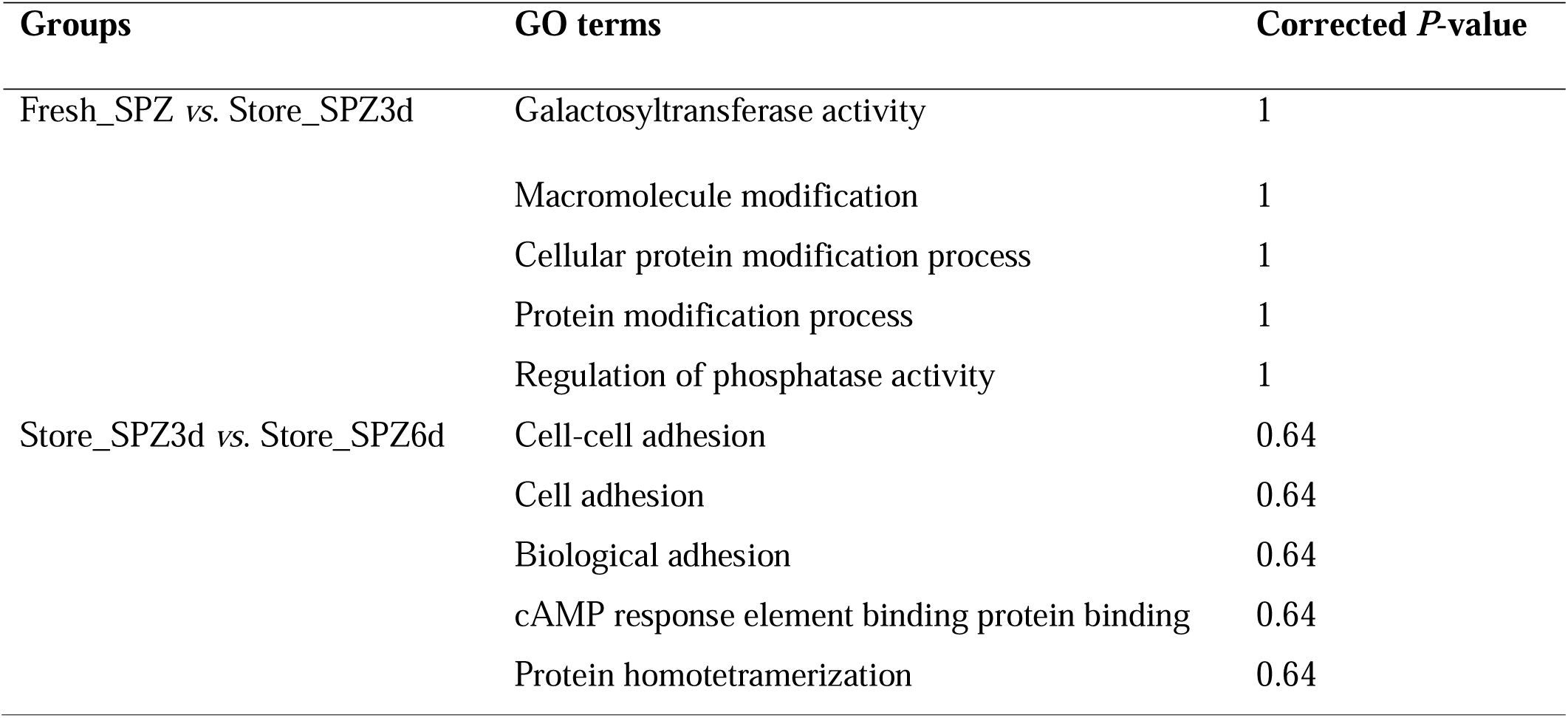
GO enrichment analysis of hypo-differentially methylated regions target genes (DMGs) in promoter (top five terms)

**Table S3.**
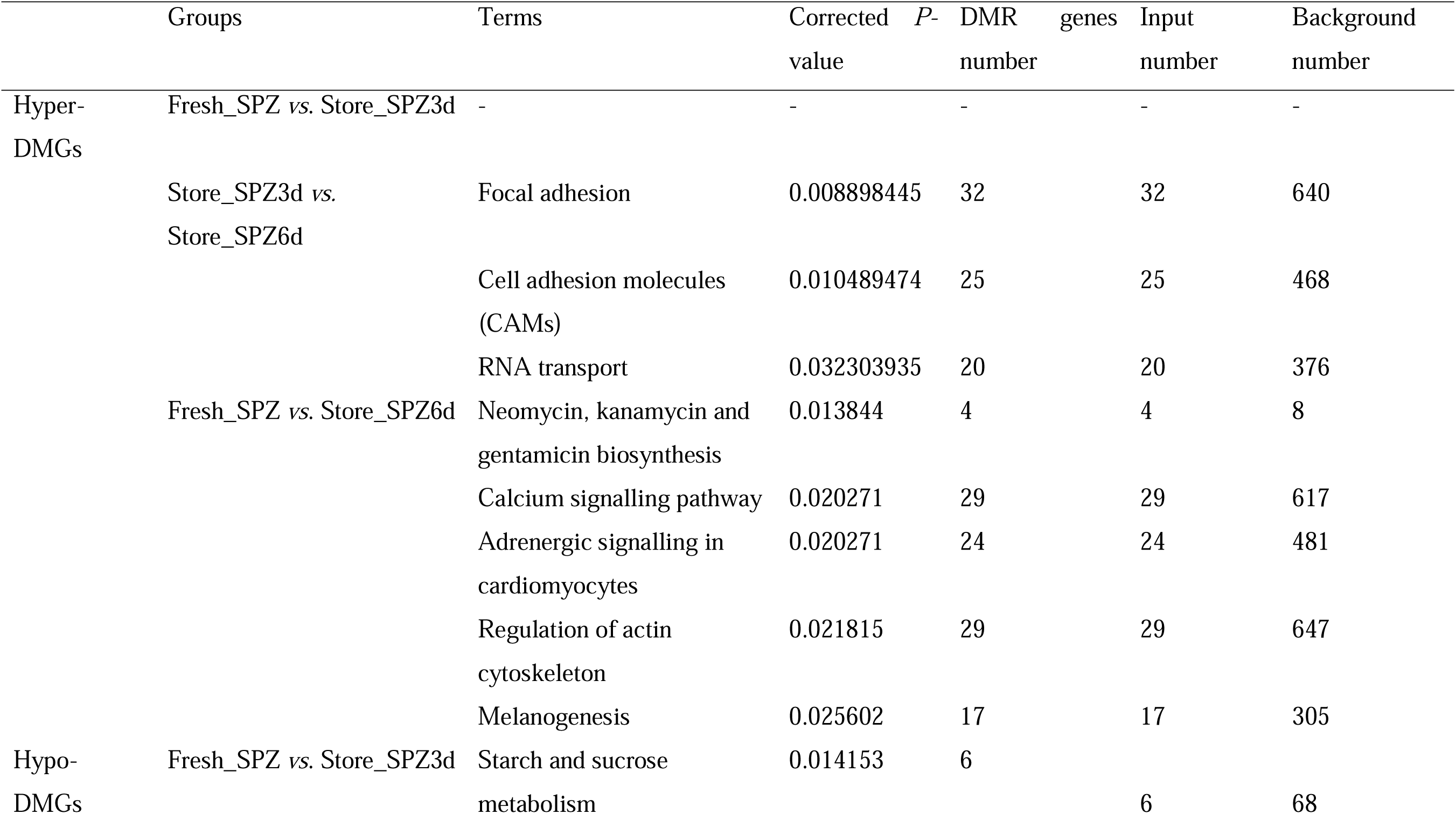
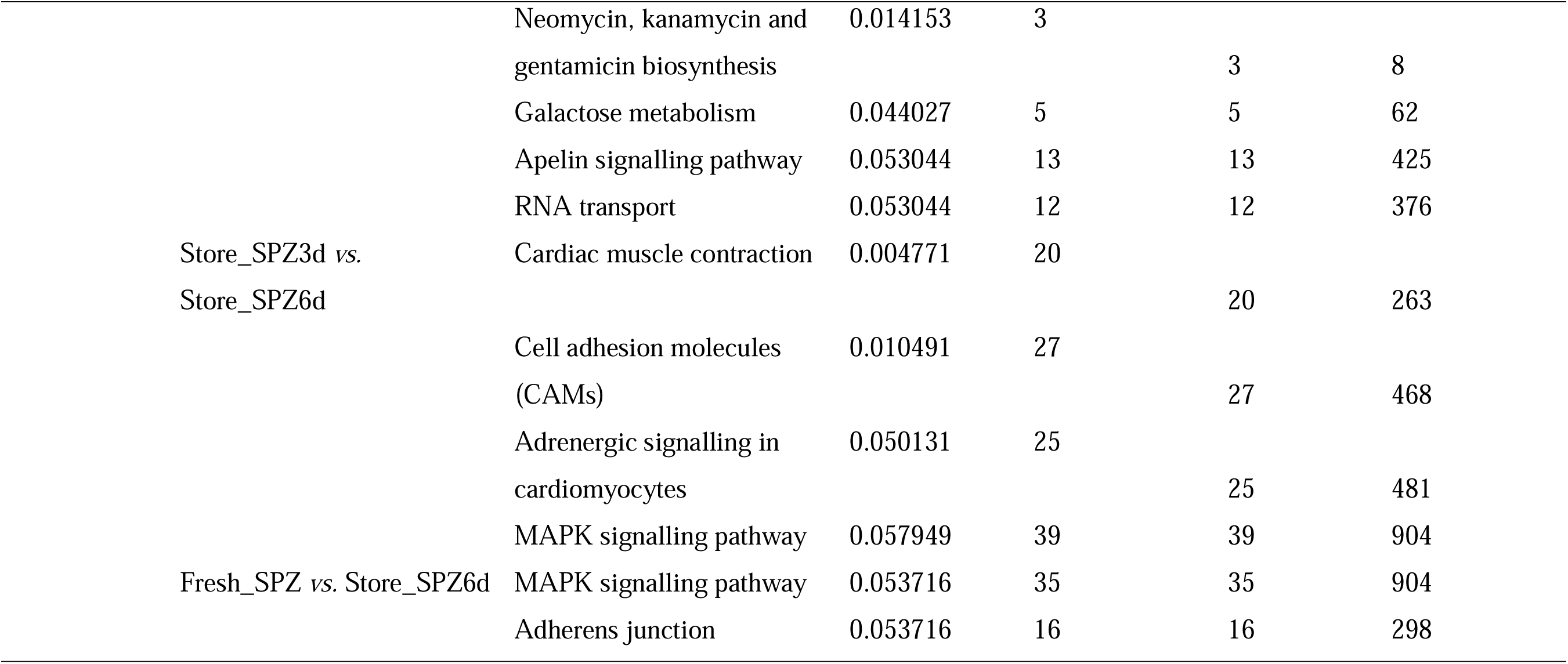
KEGG pathways significantly enriched for differentially methylated regions target genes (DMGs) in gene body (top terms with *p* ≤ 0.05)

**Fig. S1.**
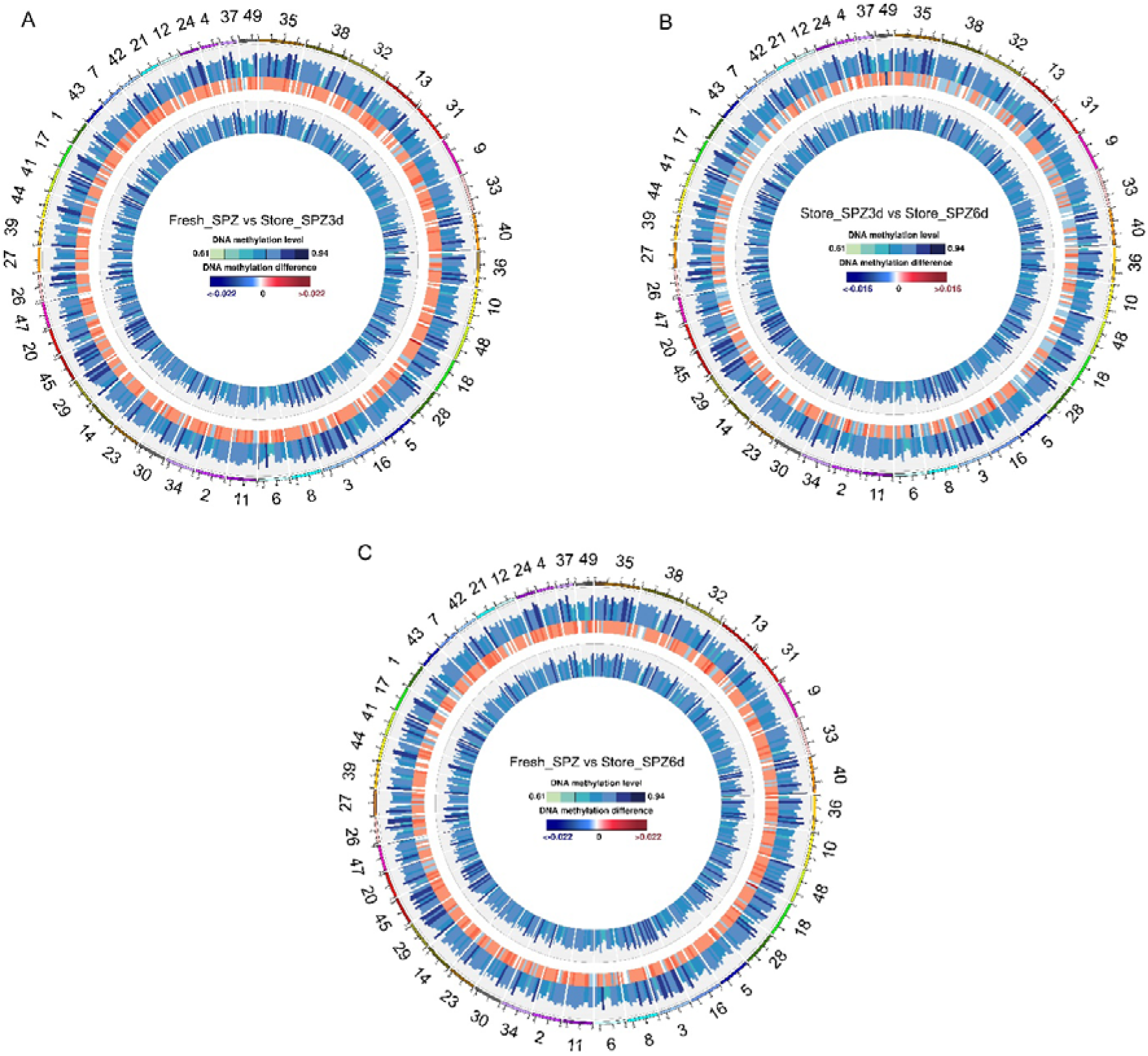
Comparison of methylation levels in CpG regions among embryos derived from fresh sperm (Fresh_SPZ), 3-day-stored (Store_SPZ3d) and 6-day-stored sperm (Store_SPZ6d). (**A**) The outer circle is embryos from fresh sperm, and the inner circle is embryos from 3-day-stored sperm. (**B**) The outer circle is embryos from 3-day-stored sperm, and the inner circle is embryos from 6-day-stored sperm. (**C**) The outer circle is embryos from fresh sperm, and the inner circle is embryos from 6-day-stored sperm. The middle circle is differences in methylation levels between two groups. The colour of the middle circle represents the difference in methylation levels. The redder the colour is, the higher methylation level of the former group than the latter group, the bluer the colour is, the lower the methylation level is in the former group than in the latter group, *n* = 3 fish replicates for each group.

**Fig. S2.**
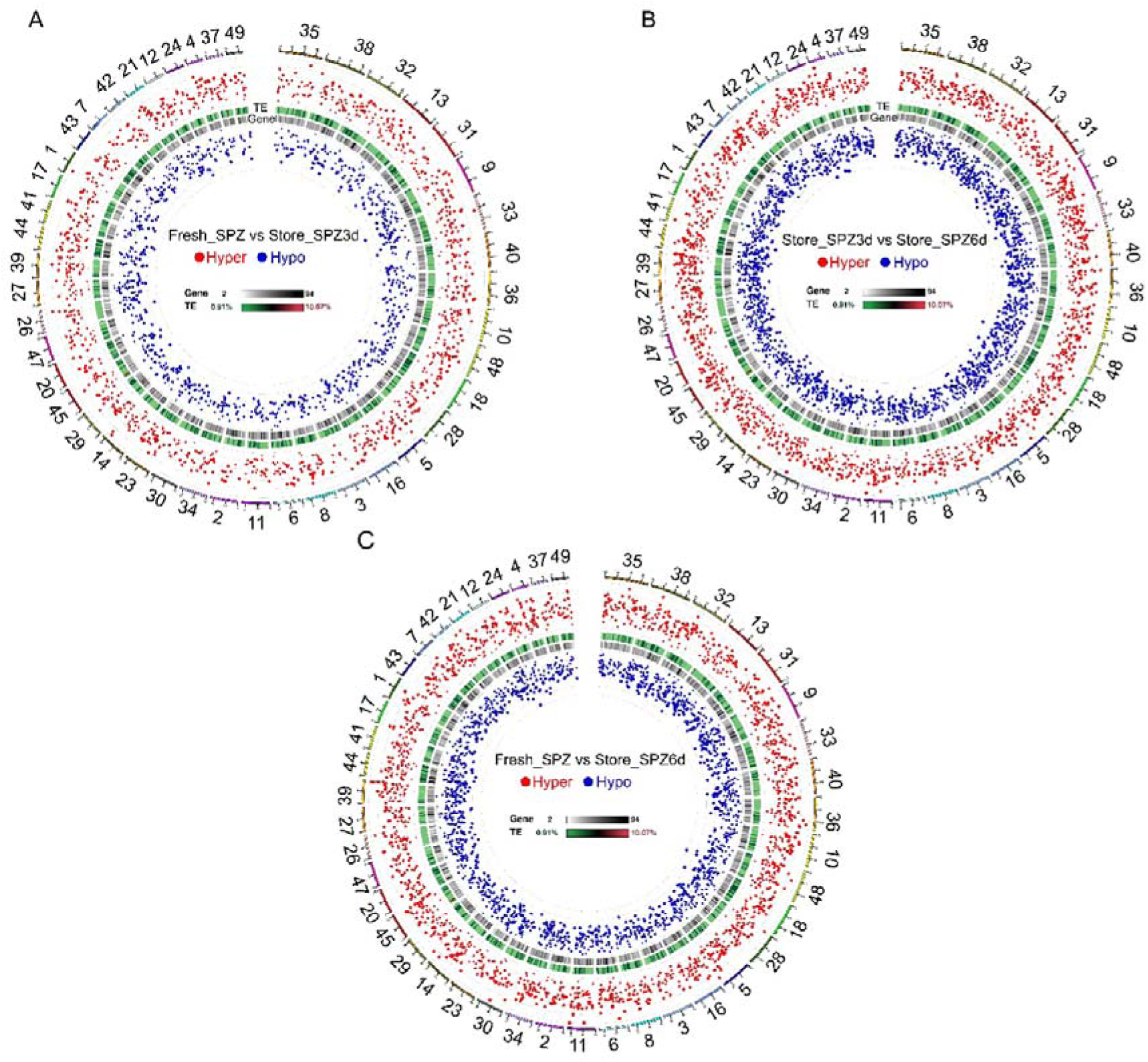
The distribution of DMRs in the genome comparison among embryos derived from fresh, 3-day-stored and 6-day-stored sperm groups. Red dots indicate hyper-DMRs, while blue dots represent hypo-DMRs.

**Fig. S3.**
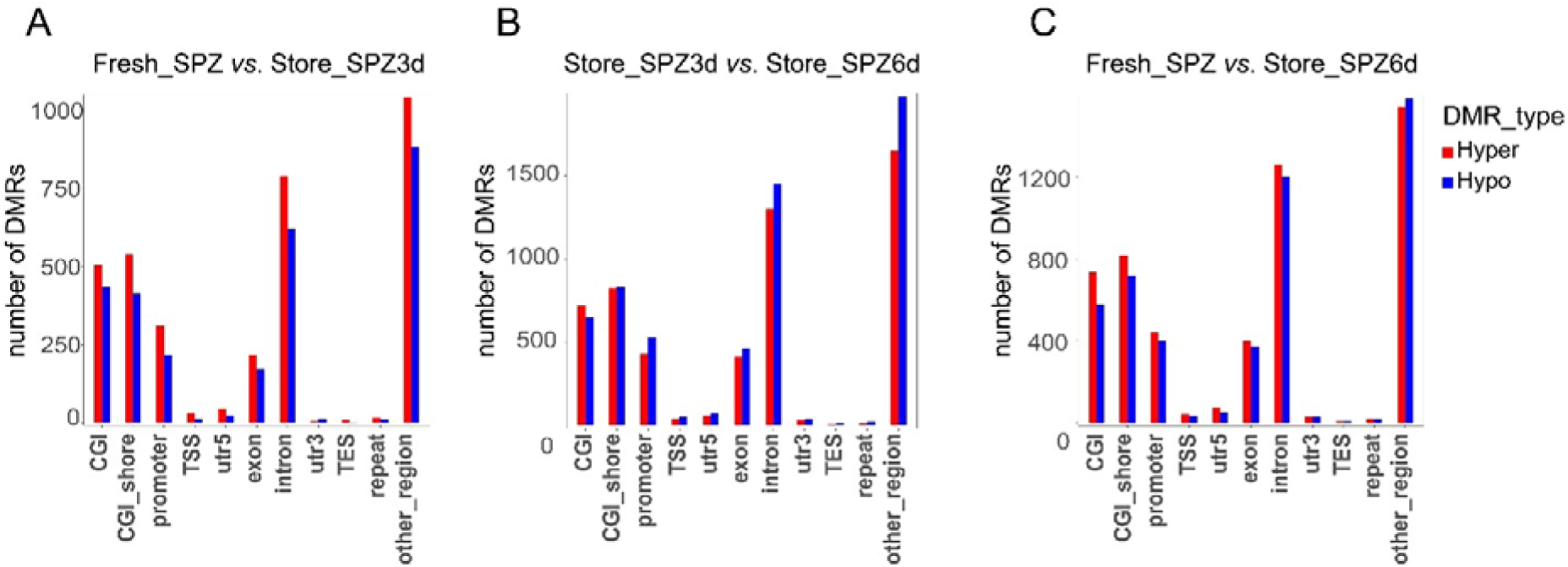
Distribution of hyper- and hypo-methylated DMRs numbers among fresh, stored sperm for 3 and 6 days *in vitro* in different gene regions. Red bar: hyper-methylated, blue bar: hypo-methylated. *n* = 3 fish replicates for Fresh_SPZ, Store_SPZ3d and Store_SPZ6d.

**Fig. S4.**
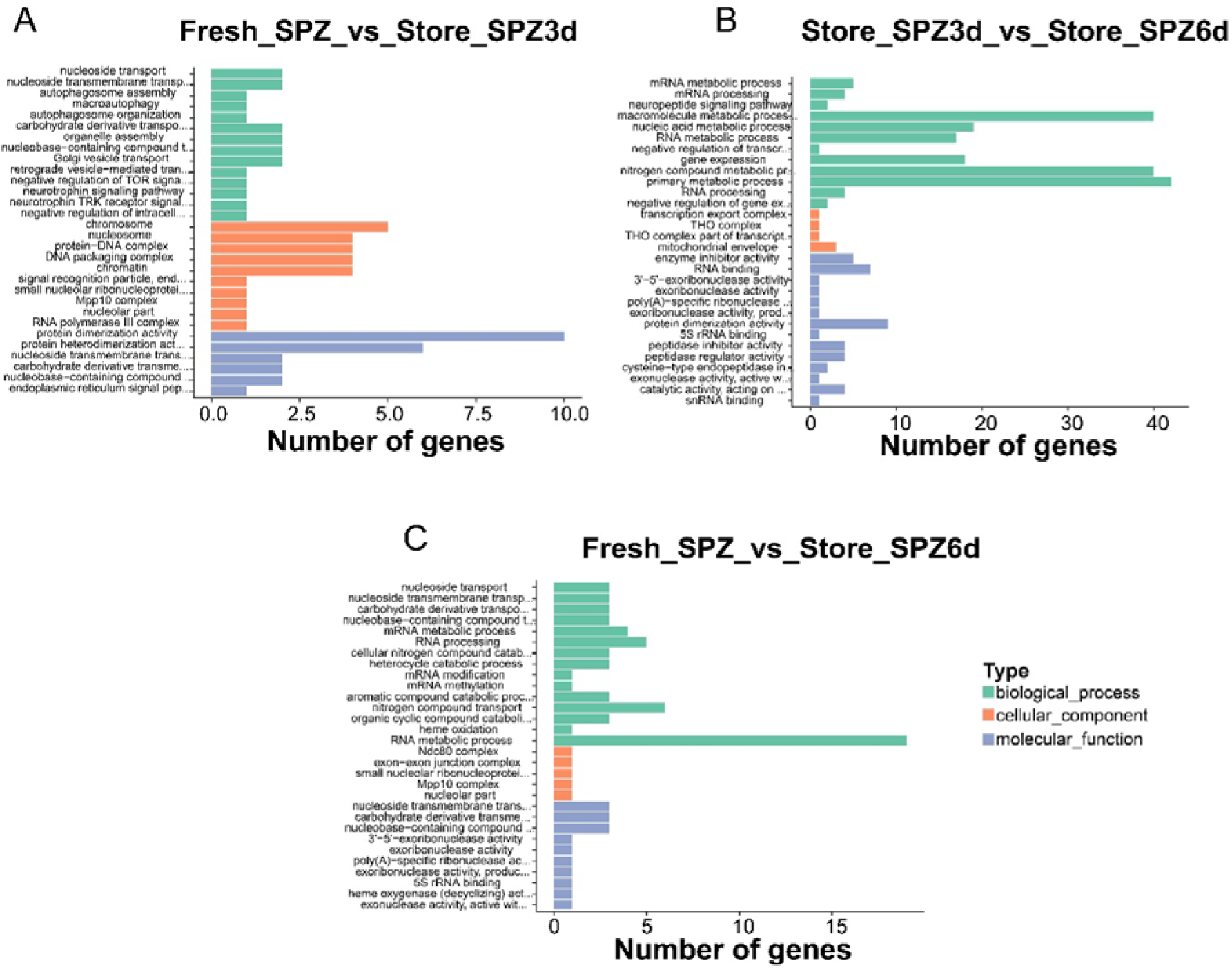
GO enrichment analysis of promoter hyper-DMGs related to CpG region

**Fig. S5.**
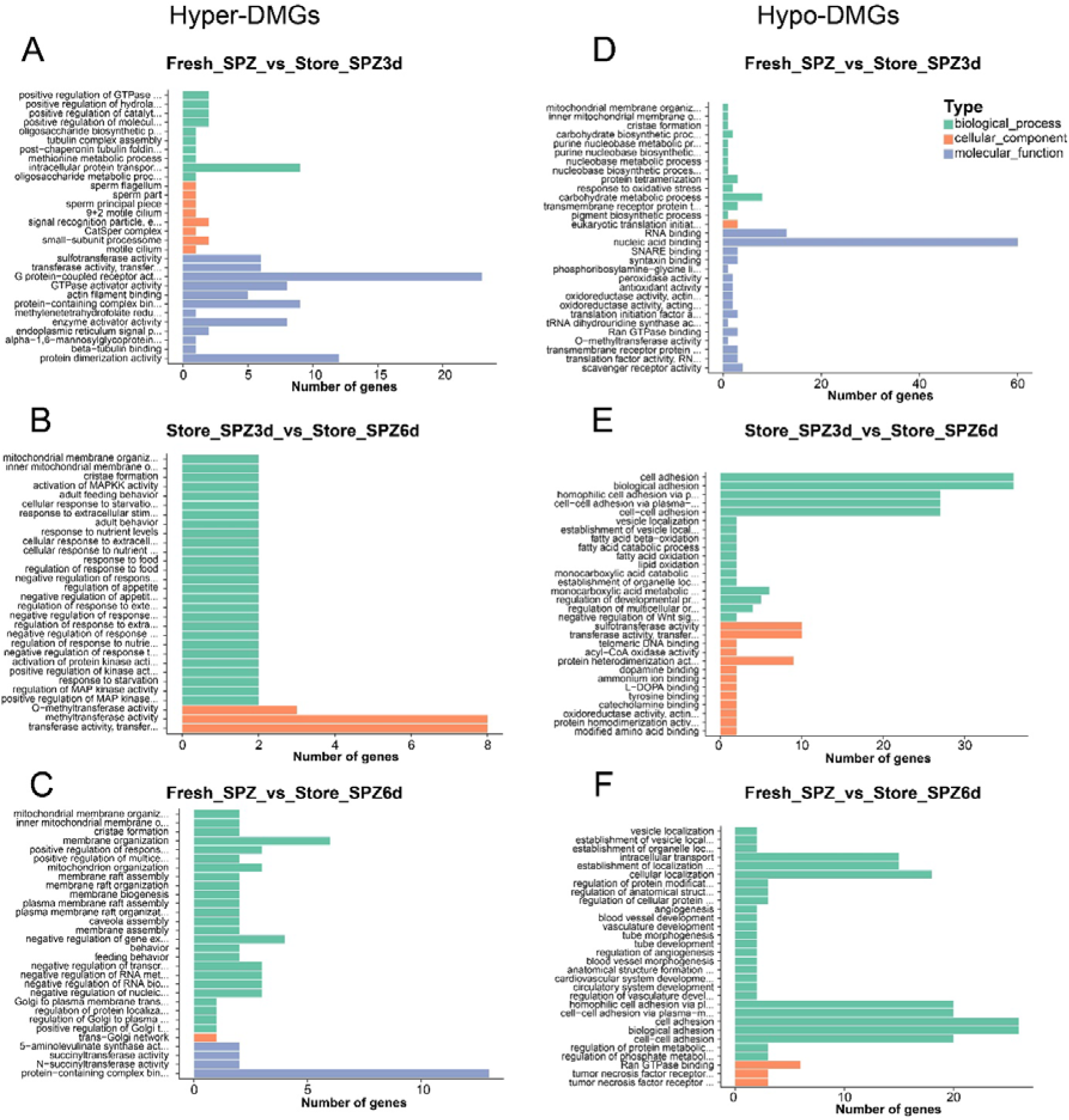
(A)-(C) GO enrichment analysis of gene body hyper-) and (D)-(F) hypo-DMGs related to the CpG region

## Notes

### Competing Interest Statement

The authors have declared no competing interest.

