## Supplementary material for "Fertilization by short-term stored sperm alters DNA methylation patterns in common carp (*Cyprinus carpio*) embryos at single-base resolution": Supplemetal Tables and Figures

**Table 1** Percentage of total genome methylation statistics. Data represent means ± S.D. (*n* = 3 fish replicates)

| Groups | Mean mC(%) | Mean mCpG(%) | Mean mCHG(%) | Mean mCHH(%) |
| --- | --- | --- | --- | --- |
| Fresh_SPZ | 7.95 ± 0.32 | 79.73 ± 0.62 | 0.93 ± 0.23 | 1.01 ± 0.26 |
| Store_SPZ3d | 7.86 ± 0.14 | 79.65 ± 0.69 | 0.78 ± 0.11 | 0.86 ± 0.13 |
| Store_SPZ6d | 7.93 ± 0.13 | 79.74 ± 0.49 | 0.90 ± 0.14 | 1.00 ± 0.17 |

No significant difference was found among three groups within each parameter (*p* > 0.05). Mean mC (%): the average methylation level of all C sites in genome, mC counts/(C counts on the genome)*100%; mean mCpG (%): average methylation levels in CpG regions, mCpG counts/(CpG counts on the genome)x100; mean mCHG (%): average methylation levels in CHG region, mCHG counts/(CHG counts on the genome) x100; mean mCHH (%): average methylation levels in CHH region, mCHH counts/(CHH counts on the genome)x100. Embryos originated from different aged sperm; fresh (Fresh_SPZ), stored for 3 days (Store_SPZ3d) and 6 days (Store_SPZ6d).

**Table 2** GO enrichment analysis of promoter hypo-DMGs (top five terms) in embryos from fresh sperm *vs*. 6-day-stored sperm

| **GO terms** | **Corrected *p*-value** | **DMR genes number** |
| --- | --- | --- |
| Cell-cell adhesion | 0.00270 | 163 |
| Homophilic cell adhesion *via* plasma membrane adhesion molecules | 0.00384 | 163 |
| Cell-cell adhesion *via* plasma-membrane adhesion molecules | 0.00384 | 163 |
| cell adhesion | 0.00384 | 163 |
| Biological adhesion | 0.00384 | 163 |

**Supplementary Files**

**Table S1** Genome coverage statistical analysis.

| Group | Total reads | Mapping rate (%) | Sites_Num | Sites_Covg | Sites_NumCovg1 | Sites_NumCovg10 |
| --- | --- | --- | --- | --- | --- | --- |
| Fresh_SPZ1 | 119,522,286 | 56.70 | 1,300,084,860 | 8.63 | 75.87% | 41.92% |
| Fresh_SPZ2 | 113,756,336 | 55.14 | 1,285,991,811 | 8.01 | 75.04% | 38.31% |
| Fresh_SPZ3 | 106,273,914 | 47.61 | 1,271,754,168 | 6.47 | 74.21% | 28.82% |
| Store_SPZ3d1 | 111,758,874 | 57.31 | 1,289,273,426 | 8.21 | 75.24% | 39.95% |
| Store_SPZ3d2 | 125,745,516 | 55.76 | 1,291,346,618 | 9.60 | 75.36% | 44.90% |
| Store_SPZ3d3 | 99,861,626 | 57.52 | 1,278,056,260 | 7.42 | 74.58% | 35.27% |
| Store_SPZ6d1 | 100,767,737 | 58.17 | 1,285,379,893 | 7.54 | 75.01% | 36.08% |
| Store_SPZ6d2 | 113,708,587 | 56.73 | 1,299,884,496 | 8.17 | 75.85% | 39.70% |
| Store_SPZ6d3 | 114,268,603 | 57.37 | 1,291,939,131 | 8.35 | 75.39% | 40.33% |

Fresh_SPZ: embryo derived from fresh sperm; Store_SPZ3d: embryo derived from stored sperm for 3 days in vitro; Store_SPZ6d: embryo derived from stored sperm for 6 days in vitro; Total reads: number of reads that have been filtered (Clean data); Mapping rate (%): ratio of reads to the total reads on the only match; sites_num: number of loci detected on the genome; Sites_CovgMean: mean coverage depth of all sites in the genome; Sites_NumCovg1: the ratio of the number of bases greater than or equal to 1× the depth of sequencing to the total length of the genome on the genome; Sites_NumCovg10: the proportion of bases greater than or equal to 10 × depth of sequencing on the genome to the total length of the genome.

**Table S2** GO enrichment analysis of hypo-differentially methylated regions target genes (DMGs) in promoter (top five terms)

| **Groups** | **GO terms** | **Corrected *P*-Value** |
| --- | --- | --- |
| Fresh_SPZ *vs.* Store_SPZ3d | Galactosyltransferase activity | 1 |
|  | Macromolecule modification | 1 |
|  | Cellular protein modification process | 1 |
|  | Protein modification process | 1 |
|  | Regulation of phosphatase activity | 1 |
| Store_SPZ3d *vs.* Store_SPZ6d | Cell-cell adhesion | 0.64 |
|  | Cell adhesion | 0.64 |
|  | Biological adhesion | 0.64 |
|  | cAMP response element binding protein binding | 0.64 |
|  | Protein homotetramerization | 0.64 |

**Table S3** KEGG pathways significantly enriched for differentially methylated regions target genes (DMGs) in gene body (top terms with *p* ≤ 0.05)

|  | **Groups** | **Terms** | **Corrected *p-* Value** | **DMR genes number** | Input number | Background number |
| --- | --- | --- | --- | --- | --- | --- |
| Hyper-DMGs | Fresh_SPZ *vs.* Store_SPZ3d | - | - | - | - | - |
|  | Store_SPZ3d *vs.* Store_SPZ6d | Focal adhesion | 0.008898445 | 32 | 32 | 640 |
|  |  | Cell adhesion molecules (CAMs) | 0.010489474 | 25 | 25 | 468 |
|  |  | RNA transport | 0.032303935 | 20 | 20 | 376 |
|  | Fresh_SPZ *vs.* tore_SPZ6d | Neomycin, kanamycin and gentamicin biosynthesis | 0.013844 | 4 | 4 | 8 |
|  |  | Calcium signalling pathway | 0.020271 | 29 | 29 | 617 |
|  |  | Adrenergic signalling in cardiomyocytes | 0.020271 | 24 | 24 | 481 |
|  |  | Regulation of actin cytoskeleton | 0.021815 | 29 | 29 | 647 |
|  |  | Melanogenesis | 0.025602 | 17 | 17 | 305 |
| Hypo-DMGs | Fresh_SPZ *vs.* tore_SPZ3d | Starch and sucrose metabolism | 0.014153 | 6 | 6 | 68 |
|  |  | Neomycin, kanamycin and gentamicin biosynthesis | 0.014153 | 3 | 3 | 8 |
|  |  | Galactose metabolism | 0.044027 | 5 | 5 | 62 |
|  |  | Apelin signalling pathway | 0.053044 | 13 | 13 | 425 |
|  |  | RNA transport | 0.053044 | 12 | 12 | 376 |
|  | Store_SPZ3d *vs.* Store_SPZ6d | Cardiac muscle contraction | 0.004771 | 20 | 20 | 263 |
|  |  | Cell adhesion molecules (CAMs) | 0.010491 | 27 | 27 | 468 |
|  |  | Adrenergic signalling in cardiomyocytes | 0.050131 | 25 | 25 | 481 |
|  |  | MAPK signalling pathway | 0.057949 | 39 | 39 | 904 |
|  | Fresh_SPZ *vs.* Store_SPZ6d | MAPK signalling pathway | 0.053716 | 35 | 35 | 904 |
|  |  | Adherens junction | 0.053716 | 16 | 16 | 298 |
